# Plasmalogen metabolism shapes germinal centre immunity

**DOI:** 10.64898/2026.03.03.706961

**Authors:** Ilaria Dorigatti, Sarah Spoeck, Denise Kummer, Katharina Lackner, Felix Eichin, Marina A. Schapfl, Georg Golderer, Verena Labi, Katrin Watschinger

## Abstract

Plasmalogens are membrane lipids whose roles in immunity are emerging through identification of the *PEDS1* gene and advances in lipid analytics. Because PEDS1 catalyses the final step of plasmalogen biosynthesis, *Peds1* knockout models plasmalogen loss. Building on reports of a haematopoietic phenotype in *Peds1*-deficient mice, we performed extended profiling to characterise haematological and immune phenotypes. In line with these previous investigations, mice showed reduced red blood cell counts and haematocrit, with increased mean platelet volume. At steady state, plasmalogen deficiency was associated with reduced germinal centre (GC) B cell frequencies in mucosal lymphoid tissues and lower basal serum IgG1 levels. Following immunisation, GC B cell frequencies and ovalbumin-specific IgG1 were similar across genotypes, yet GC B cells exhibited elevated mitochondrial superoxide and plasma cells displayed increased lipid peroxidation. To test for B cell-intrinsic effects, we used an *in vitro* GC culture, where *Peds1*-deficient B cells showed reduced class switching to IgG1, intracellular IgG1 accumulation and diminished IgG1 secretion. Together, these findings support a role for plasmalogens and PEDS1 as a lipid-metabolic modulator of GC dynamics and IgG1 output, consistent with broader ether lipid perturbations. Plasmalogen metabolism may therefore offer a potential entry point to modulate humoral immunity or haematopoiesis.

## Introduction

The mammalian lipidome comprises thousands of species that support membrane compartmentalisation, signalling, and energy metabolism. These lipids are organised into a variety of categories, classes and subclasses, as depicted in the LIPID MAPS database^1,2^. Ether lipids are defined by an ether bond at the *sn*-1 position of the glycerol backbone and occur in both glycerophospholipid and neutral glycerolipid classes. They have been increasingly investigated over the last decades and advances in mass spectrometry now allow much deeper coverage of these lipid species in complex lipid mixtures^3,4^. Ether lipid biosynthesis initiates in peroxisomes and continues at the endoplasmic reticulum (ER)^5^. The first peroxisomal steps include acylation of glyceronephosphate by the enzyme glyceronephophate-*O*-acyltransferase (GNPAT, EC: 2.3.1.42), subsequent exchange of this acyl side chain by the enzyme alkylglycerone phosphate synthase (AGPS, EC: 2.5.1.26), using a fatty alcohol provided by the rate-limiting enzymes fatty acid reductase 1 and 2 (FAR1/2, EC: 1.2.1.84). The reduction of the *sn*-2 keto group is then accomplished by acyl/alkylglycerone phosphate reductase (DHRS7B, EC: 1.1.1.101), which resides both at peroxisomal level but also at the ER, where all further maturation steps take place^5^. Ether lipids comprise saturated alkyl ether lipids (also called plasmanyl lipids) and alkenyl ether lipids, better known as plasmalogens, defined by a vinyl ether double bond. The introduction of this distinguishing double bond in plasmalogens is catalysed by the ER-resident enzyme plasmanylethanolamine desaturase (PEDS1, EC: 1.14.19.77), whose gene was recently identified (gene symbol: *PEDS1*, former gene symbol: *TMEM189*)^6–8^.

Human genetic disorders underscore the physiological relevance of ether lipids. Disruptions in peroxisomal ether lipid biosynthesis cause Rhizomelic Chondrodysplasia Punctata (RCDP) while global peroxisomal dysfunctions lead to diseases of the Zellweger spectrum^9,10^. In both disorders, ether lipid biosynthesis is profoundly impaired, causing severe neurological, skeletal and ocular abnormalities, often culminating in death within the first decade of life^9–11^. Case reports of patients with Zellweger syndrome and related peroxisome biogenesis disorders document severe infections, haematologic abnormalities such as anaemia and thrombocytopenia, and reduced erythrocyte ethanolamine plasmalogen levels in early infancy^10,12^. In addition, complete absence of B cells, markedly reduced immunoglobulin (Ig) levels, and poor IgG increase despite replacement therapy were described in an infant with genetically confirmed Zellweger syndrome, consistent with a defect in B cell generation and antibody-mediated responses to infection^13^. Human *PEDS1* variants causing selective plasmalogen deficiency were unknown until 2026, when Amelan A. *et al.* first reported two male infants harbouring biallelic *PEDS1* mutations^14^. These patients presented with microcephaly, developmental delay, impaired neurogenesis, recurrent chest infections suggestive of immune dysfunction, and unilateral cataract in one case.

Ether lipids including plasmalogens are widely present in many tissues and membranes of the body. They make up a significant portion of immune cell membranes^15^ and can modulate the immune response, highlighting their pivotal role in immunity^16^. Mechanistic work links ether lipids to both innate and adaptive immunity. In a *peds1*-deficient zebrafish model, selective plasmalogen loss caused myeloid cell apoptosis and exacerbated inflammation^17^. A dependence of macrophages on ethanolamine plasmalogen was described in investigations on macrophage function conducted in murine RAW.108 cells^18^, an ether lipid-deficient macrophage cell line, and bone marrow-derived macrophages (BMDMs) from *Lpcat3/Elovl5* double KO mice, which significantly reduces polyunsaturated fatty acid (PUFA) phospholipids content^19^. These lipids are highly abundant and regulate membrane properties, such as fluidity and lipid-rich microdomain formation, which are crucial for phagocytosis^18^. Plasmalogens also shift under metabolic stress in human monocyte-derived macrophages^20^, linking ether lipid metabolism directly to immunometabolism. Neutrophil studies in two inducible KO mouse models - for *Fas* (fatty acid synthase, catalysing *de novo* lipogenesis) and *Dhrs7b* - demonstrated that acute ether lipid loss is associated with neutropenia via a shift in membrane phospholipid composition, and impaired cell viability and tissue recruitment during induced inflammation^21,22^. However, a parallel research reported no systemic neutropenia or leukopenia in chronic ether lipid deficiency in mice lacking *Gnpat*, the first peroxisomal enzyme GNPAT of ether lipid biosynthesis, and human patients suffering from RCDP, suggesting a potential compensatory mechanism in a sustained deficiency state^23^. 1-*O*-Alkyl-2-acetyl-*sn*-glycero-3-phosphocholine, known as platelet-activating factor (PAF), a potent pro-inflammatory ether lipid, is produced by various innate immune cells (neutrophils, eosinophils, macrophages and platelets, but also endothelial cells)^24^ in response to inflammatory stimuli, such as phagocytosis, or oxidative stress, like reactive oxygen species (ROS) burst^25^.

Plasmalogens also intersect with adaptive immunity^26–28^. In the *Gnpat* KO mouse model, ether lipids act as self-antigens for invariant Natural Killer T cells (iNKT) selection in the thymus^29^. Moreover, the bone marrow, the site of haematopoiesis, contains ethanolamine plasmalogen, which account for approximately 8% of its total phospholipid pool^30^, indicating that plasmalogens are substantial components of the haematopoietic membrane lipidome. An investigation of the *Cd36* KO mouse model highlights that remodelling of phospholipids, such as phosphatidylserine, supports megakaryopoiesis and platelet production, raising the intriguing possibility that ether lipids, which modulate membrane properties in other hematopoietic lineages^28,31^, may similarly contribute to these processes^32^. A recent study in a total, B cell-specific or germinal centre (GC) B cell-specific inducible deletion of *Dhrs7b* in mouse, demonstrated that B cell-intrinsic ether lipid biosynthesis is essential for proper GC formation, antibody maturation and B cell survival. Notably, these data suggest through redox regulation a pivotal role in adaptive immunity for ether lipids^28^.

Although these studies address the consequences of global ether lipid loss in various immune cell types, such as neutrophils, macrophages, iNKT cells and B cells, we still lack a comprehensive understanding of how a selective plasmalogen deficiency affects the immune system in mammals. Such plasmalogen deficits might be therapeutically targetable via supplementation; e.g. shark liver oil is a natural ether lipid-rich source, which boosts endogenous plasmalogen levels and supports immune function while reducing inflammation in metabolic conditions^33^.

The recent identification of the *PEDS1* gene and the subsequent availability of a *Peds1*-deficient mouse model now allow *in vivo* distinction between alkyl ether lipids and plasmalogens. By phenotyping the mouse model lacking *Peds1*, we could reveal a strong reduction of plasmalogens in all tested organs and significantly decreased body weight compared to wildtype (WT) controls, underscoring the essential role of the vinyl ether double bond for normal growth^7^. Further, in lipidomic studies, we demonstrated that *Peds1* deficiency triggers compensatory plasmanyl ether lipid accumulation across the tissues, maintaining constant levels of the class of phosphatidylethanolamines^34^. Data from the International Mouse Phenotyping Consortium (IMPC) further indicate haematological abnormalities, including increased mean platelet and corpuscular volumes, alongside decreased haemoglobin content, haematocrit, and erythrocyte numbers in murine *Peds1* deficiency^35^.

To determine how selective plasmalogen deficiency affects the immune system, we here provide an integrated immunological investigation of the *Peds1* KO mouse. Our comprehensive analysis reveals that *Peds1* deficiency alters blood cell parameters and humoral immunity. Understanding plasmalogen contributions to B cell activation and GC responses is vital for further elucidating lipid regulation of adaptive immunity, paving the way for novel therapeutic concepts in immune disorders and ether lipid-related pathologies.

## Results

### *Peds1* knockout in mice affects red blood cell count and haematocrit

The IMPC reported reduced red blood cell count and haematocrit, as well as increased mean platelet volume in germline *Peds1* KO mice^35^. Prompted by these findings, we used the same model to extend this baseline phenotyping with broader haematological and immunophenotyping analyses to characterise the impact of plasmalogen loss. Despite significantly reduced body weight in 8-weeks-old *Peds1*-deficient mice (*Peds1* KO = 21.4 ± 2.79 g, WT = 24.3 ± 3.75 g, *p* = 0.022), the ratios of spleen/body weight and thymus/body weight, as well as cellularity of spleen (SPL), bone marrow (BM), mesenteric lymph nodes (LNmes), Peyer’s patches (PP), and peritoneal cavity lavage (PEC) were comparable between genotypes (Suppl. Fig. 1a-h).

In the bone marrow, we quantified long-term haematopoietic stem cells (LT-HSCs), short-term HSCs (ST-HSCs), and the broader multipotent progenitors (MPP) and LSK (Lin^-^Sca1^+^Kit^+^) compartments (Fig. 1a). These populations were comparable between *Peds1* KO and WT control littermates (Fig. 1b-e). Analysis of bone marrow-derived erythroblasts and total Ter119^+^ erythroid cells also revealed no genotype-dependent variation (Fig. 1f, g). The distribution of erythroid subpopulations A, B, and C within Ter119^+^CD71^+^ cells was also unchanged (Fig. 1h), consistent with unaltered bone marrow erythropoiesis.

**Figure 1:**
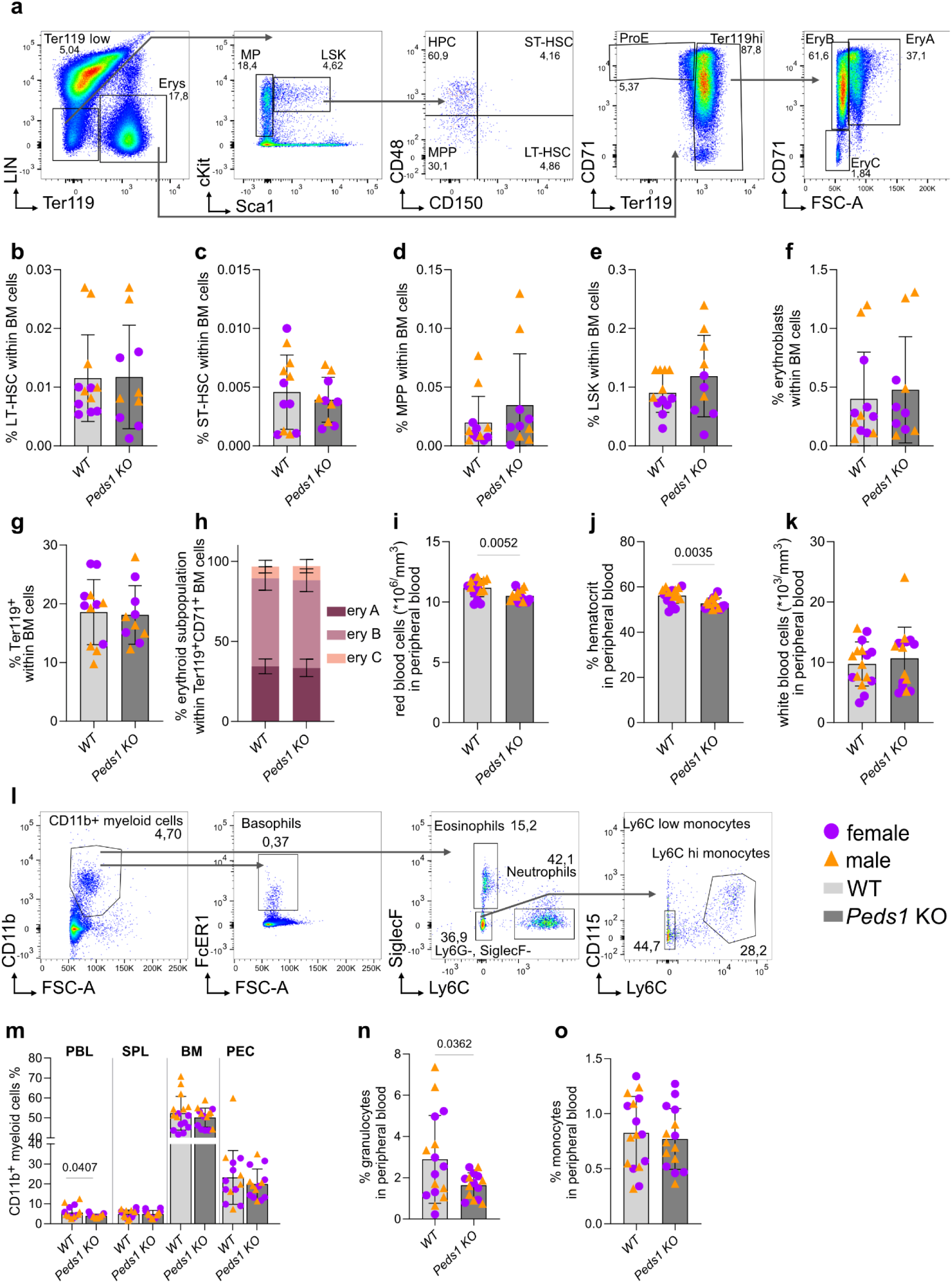
Haematological analysis and flow cytometric profiling of haematopoietic and myeloid compartments in WT and Peds1 KO mice. **a** Gating strategy for HSCs in bone marrow. **b** Long-term haematopoietic stem cells (LT-HSCs: Lin^-^, cKit^+^, Sca1^-^, CD48^-^, CD150^+^), **c** short-term haematopoietic stem cells (ST-HSC: CD48^+^, CD150^+^), **d** multipotent progenitors (MPP, CD48^-^, CD150^-^), **e** LSK cells (Lin^-^, cKit^+^, Sca1^-^), **f** erythroblasts (CD71^+^, Ter119^+^), **g** Ter119^+^ cells, **h** erythrocyte subtypes within Ter119^+^CD71^+^ cells (ery A: Ter119^+^, CD71^high^; ery B: Ter119^+^, CD71^+^; ery C: Ter119^+^, CD71^-^) in BM. **i** Red blood cell count, **j** haematocrit, and **k** white blood cell count in PBL. **l** Gating strategy for myeloid cell populations in the BM. **m** Fraction of CD11b^+^ myeloid cells in PBL, SPL, BM, and PEC. **n** Fraction of granulocytes (total of basophils: CD11b^+^, FcεR1^+^; eosinophils: CD11b^+^, SiglecF^+^; neutrophils: CD11b^+^, Ly6G^+^) in PBL, **o** fraction of monocytes (CD11b^+^, Ly6C^+^) in PBL. **b-h** WT (n = 12, 6 female, 6 male mice), Peds1 KO mice (n = 10, 5 female, 5 male mice); **i-k** WT (n = 16, 8 female, 8 male mice), Peds1 KO mice (n = 15, 8 female, 7 male mice); **m-o** WT (n = 16, 8 female, 8 male mice), Peds1 KO mice (n = 15, 8 female, 7 male mice), except for both WT and Peds1 KO in PEC (n = 14, 8 female, 6 male mice). Females are shown as violet circles and males as yellow triangles. Data are presented as mean ± SD and statistical significance was determined by two-tailed Student’s t-test (**b-k, n-o**) or one-way ANOVA with Bonferroni’s multiple comparison test (**m**).

Similar to the findings of the IMPC, peripheral blood (PBL) analyses demonstrated a marked reduction in red blood cell counts (*Peds1* KO = 10.5 ± 0.475, WT = 11.2 ± 0.743, *p* = 0.005, Fig. 1i) and haematocrit in *Peds1* KO mice compared to controls (*Peds1* KO = 52.7 ± 2.45, WT = 56.2 ± 3.44, *p* = 0.0035, Fig. 1j), whereas total white blood cell counts were unchanged (*Peds1* KO = 10.7 ± 5.19, WT = 9.74 ± 3.66, *p* = 0.569, Fig. 1k). Additional haematological parameters, including platelet count, mean corpuscular volume and haemoglobin, haemoglobin concentration, red cell distribution width, and haemoglobin levels did not differ between genotypes (Suppl. Fig. 1i, k-o). Platelet volume, however, was elevated in *Peds1* knockout mice (*Peds1* KO = 5.85 ± 0.466, WT = 5.22 ± 0.446, *p* = 0.0006, Suppl. Fig. 1j). These data largely confirmed the observations of the basic IMPC phenotyping. Next, we profiled myeloid cells (Fig. 1l). We observed a decrease in circulating CD11b^+^ myeloid cell frequency in *Peds1* knockout animals (PBL: *Peds1* KO = 3.39 ± 0.92, WT = 5.66 ± 2.99, *p* = 0.0407, Fig. 1m), while frequencies in spleen, bone marrow, and peritoneal cavity lavage were unchanged (Fig. 1m). In peripheral blood, granulocyte frequencies were reduced (Fig. 1n), while monocyte frequencies were unvaried (Fig. 1o). Further, no genotype-dependent differences were detected in eosinophils, granulocytes, monocytes, or lymphocytes in peripheral blood (Suppl. Fig. 1p). Neutrophils, basophils, eosinophils, Ly6C high and Ly6C low monocytes were assessed by flow cytometry in peripheral blood (Suppl. Fig. 2a-e), in spleen (Suppl. Fig. 2f-j), in bone marrow (Suppl. Fig. 2k-o) and peritoneal cavity lavage (Suppl. Fig. 2p-t). Significant differences were detected only in granulocytes (Fig. 1n), neutrophils (Suppl. Fig. 2a) and eosinophils (Suppl. Fig. 2c) in peripheral blood of *Peds1-*deficient mice.

In conclusion, *Peds1*-deficient mice present with significant changes in haematopoietic parameters, an increase in mean platelet volume and no significant alterations in white blood cell and platelet counts. Alterations observed in the myeloid compartment were confined specifically to granulocytes, neutrophils and eosinophils in peripheral blood. Throughout our analyses, no sex-specific phenotypes were identified. Given that our data recapitulated the publicly available data from the IMPC^35^ we proceeded to an in depth analysis of selective plasmalogen deficiency beyond what was published by the IMPC.

### *Peds1* deficiency does not perturb T cell development but is associated with reduced circulating CD8^+^ T cells

Building on the reported role of plasmalogens in modulating T cell membrane composition and signaling^29^, we examined how *Peds1* deficiency influences T cell homeostasis and function in our knockout model under steady state conditions. We first assessed thymic T cell development by flow cytometry (Fig. 2a, Suppl. Fig. 3a). Overall, thymocyte subset distribution across the major developmental stages was similar between genotypes (Suppl. Fig. 3b-l). One exception was a mild increased frequency of early T cell progenitors (ETP) in *Peds1* KO mice (*Peds1* KO = 0.013 ± 0.002, WT = 0.008 ± 0.002, *p* = 0.012, Suppl. Fig. 3e).

**Figure 2:**
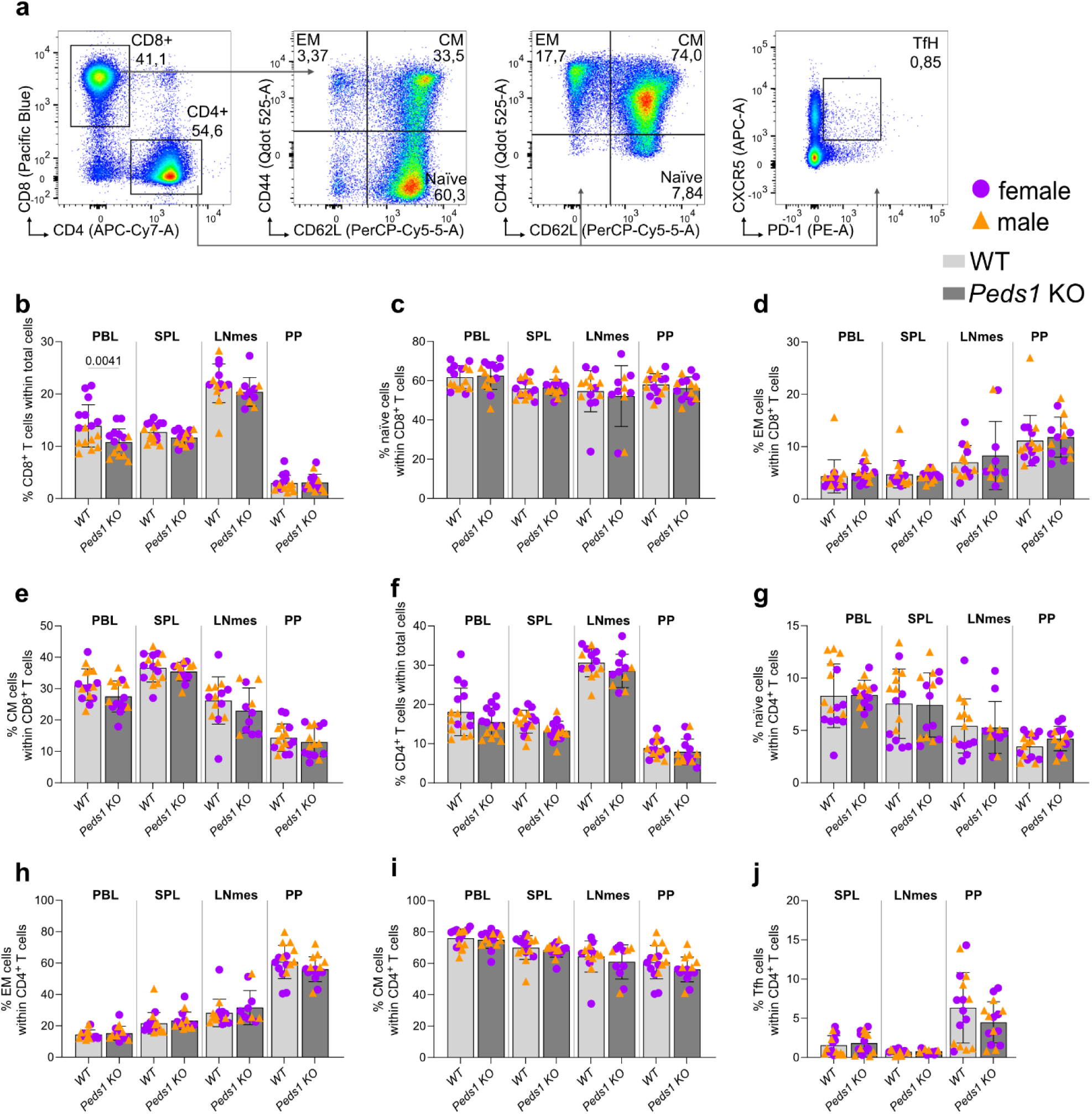
T cell subset distribution in peripheral lymphoid tissues of WT and Peds1 KO mice. Flow cytometric analysis of T cell populations in WT (n = 16, 8 female, 8 male mice; in LNmes: n = 14, 7 female, 7 male) versus Peds1 KO mice (n = 11, 7 female, 4 male mice). All cell populations were reported in four different tissues: PBL, SPL, LNmes and PP. **a** Gating strategy for T cell subsets. **b** Fraction of CD8^+^ T cells. **c** Fraction of naïve CD8^+^ T cells (CD44^-^, CD62L^+^). **d** Fraction of CD8^+^ effector memory T cells (EM, CD44^+^, CD62L^-^). **e** Fraction of CD8^+^ central memory T cells (CM, CD44^+^, CD62L^+^). **f** Fraction of CD4^+^ T cells. **g** Fraction of naïve CD4^+^ T cells (CD44^-^, CD62L^+^). **h** Fraction of CD4^+^ effector memory T cells (EM, CD44^+^, CD62L^-^). **i** Percentage of CD4^+^ central memory T cells (CM, CD44^+^, CD62L^+^). **j** Fraction of T follicular helper CD4^+^ T cells (Tfh, CXCR5^+^, PD-1^+^). Females are shown as violet circles and males as yellow triangles. Data are presented as mean ± SD. Statistical significance was determined by one-way ANOVA with Bonferroni’s multiple comparison test.

We next quantified peripheral T cell subsets in peripheral blood and secondary lymphoid organs. Circulating CD8^+^ T cells were reduced in peripheral blood of *Peds1* KO mice (*Peds1* KO = 10.8 ± 2.55, WT = 13.9 ± 4.05, *p* = 0.016), whereas no alterations were observed in the spleen, mesenteric lymph nodes, or Peyer’s patches (Fig. 2b). Stratification into naïve, effector memory (EM) and central memory (CM) CD8^+^ T cell subsets revealed no further genotype-specific differences in all tested organs (Fig. 2c-e). CD4^+^ T cells were likewise unchanged across peripheral blood, spleen, mesenteric lymph nodes, and Peyer’s patches, and their naïve, EM, CM subsets were similar between genotypes (Fig. 2f-i). In Peyer’s patches we noted a trend towards fewer CD4^+^ T follicular helper (Tfh) cells (*Peds1* KO = 4.48 ± 2.62, WT = 6.25 ± 4.50, *p* = 0.173, Fig. 2j), a T cell subset which is essential for GC formation and provides crucial help to B cells during the germinal centre (GC) reaction^36^.

Finally, NK and NKT cell frequencies were comparable across peripheral blood, spleen, mesenteric lymph nodes, Peyer’s patches and peritoneal cavity lavage (Suppl. Fig. 3m-o). Given the Tfh trend in Peyer’s patches indicating a potential alteration in GC-associated immunity, we next examined B cell populations in more detail.

### Plasmalogen deficiency reduces GC B cell frequency and IgG1 antibody production at steady state

Recent work by Cho *et al.* reported enrichment of ether lipids in GC B cells and showed that loss of DHRS7B, an enzyme upstream of PEDS1 in ether lipid biosynthesis, perturbs B cell homeostasis and proliferation with smaller GC structures and impaired antibody responses^28^. To test the specific contribution of plasmalogen deficiency *in vivo*, we analysed the B cell compartment in *Peds1*-deficient mice, a setting in which plasmanyl lipids are retained.

We first quantified total B cells across tissues. The B220^+^CD19^+^ B cell fraction was increased in the peripheral blood of *Peds1* KO mice, while frequencies in spleen, lymph nodes, Peyer’s patches and peritoneal cavity lavage were unvaried (Fig. 3a and b). Analysis of B cell development in the bone marrow and final maturation in the spleen revealed no substantial genotype-dependent differences (Suppl. Fig. 4a-o).

**Figure 3:**
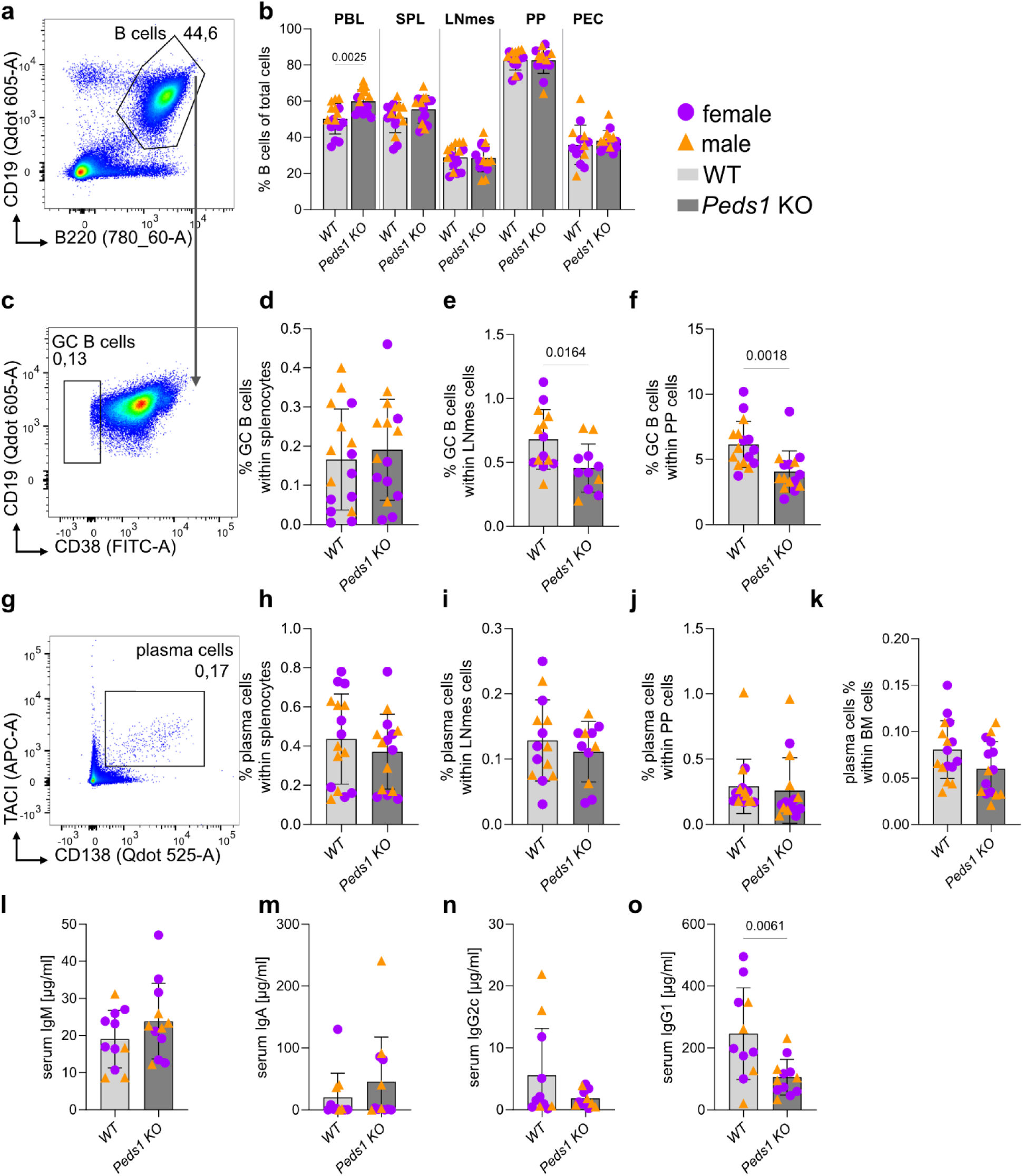
B cell compartment analysis and serum immunoglobulins in WT and Peds1 KO mice. Flow cytometric analyses of B cell populations and ELISA-based quantification of serum Igs in WT (n = 16, 8 female, 8 male mice; in LNmes and serum: n = 14, 7 female, 7 male) versus Peds1 KO mice (n = 15, 8 female, 7 male mice; in LNmes: n = 11, 7 female, 4 male; in serum: n = 12, 7 female, 5 male). Analyses were performed from PBL, SPL, LNmes, PP, and PEC. **a** Gating strategy for B cells. **b** Frequency of total B cells in PBL, SPL, LNmes, PP, and PEC (B220^+^, CD19^+^). **c** Gating strategy for GC B cells (B220^+^, CD19^+^, CD38^-^), measured in **d** splenocytes, **e** LNmes, and **f** PP. **g** Gating strategy for plasma cells. **h, i, j, k** Frequency of plasma cells in SPL, LNmes, PP and BM respectively (TACI^+^, CD138^+^). **l** IgM, **m** IgA, **n** IgG2c, and **o** IgG1 levels in serum, measured via ELISA. Females are shown as violet circles and males as yellow triangles. Data are presented as mean ± SD. Statistical significance for B cell percentages in **b** was determined using a one-way ANOVA with Bonferroni’s multiple comparisons test. All other comparisons were performed using a two-tailed unpaired Student’s t-test.

We next focused on GC B cells. GCs are transient structures that form in secondary lymphoid tissues following antigen engagement via the B cell receptor, and support clonal expansion and differentiation towards memory B cells or antibody-secreting plasma cells^37^. In *Peds1* KO mice at steady state, GC B cell frequencies were unchanged in spleen (*Peds1* KO = 0.191 ± 0.129, WT = 0.166 ± 0.129, *p* = 0.594, Fig. 3c, d), but were significantly reduced in mesenteric lymph nodes (*Peds1* KO = 0.456 ± 0.188, WT = 0.681 ± 0.233, *p* = 0.0164, Fig. 3e) and Peyer’s patches (*Peds1* KO = 4.06 ± 1.59, WT = 6.15 ± 1.77, *p* = 0.0018, Fig. 3f). Plasma cell frequencies were comparable between genotypes in various tissues (Fig. 3g-k). Consistent with this, serum immunoglobulin levels of IgM, IgA, and IgG2c were unchanged (Fig. 3l-n). In contrast, serum IgG1 titres were significantly reduced in *Peds1* KO mice (Fig. 3o), motivating us to further examine GC-associated B cell responses.

### Plasmalogen deficiency reduces class switch recombination (CSR) and IgG1 secretion in an *in vitro* induced germinal centre (iGC) culture

To substantiate the reduction in GC B cell frequencies and lower IgG1 observed in *Peds1* knockout mice, we used an *in vitro* induced germinal centre (iGC) culture system. Naïve B cells were co-cultured with CD40L- and BAFF-expressing feeder cells in the presence of IL-4 for 4 days and IL-21 for 4 more days, to mimic key GC signals (Fig. 4a)^38^. This setup allowed us to phenotypically assess plasmalogen dependence in GC B cell biology.

**Figure 4:**
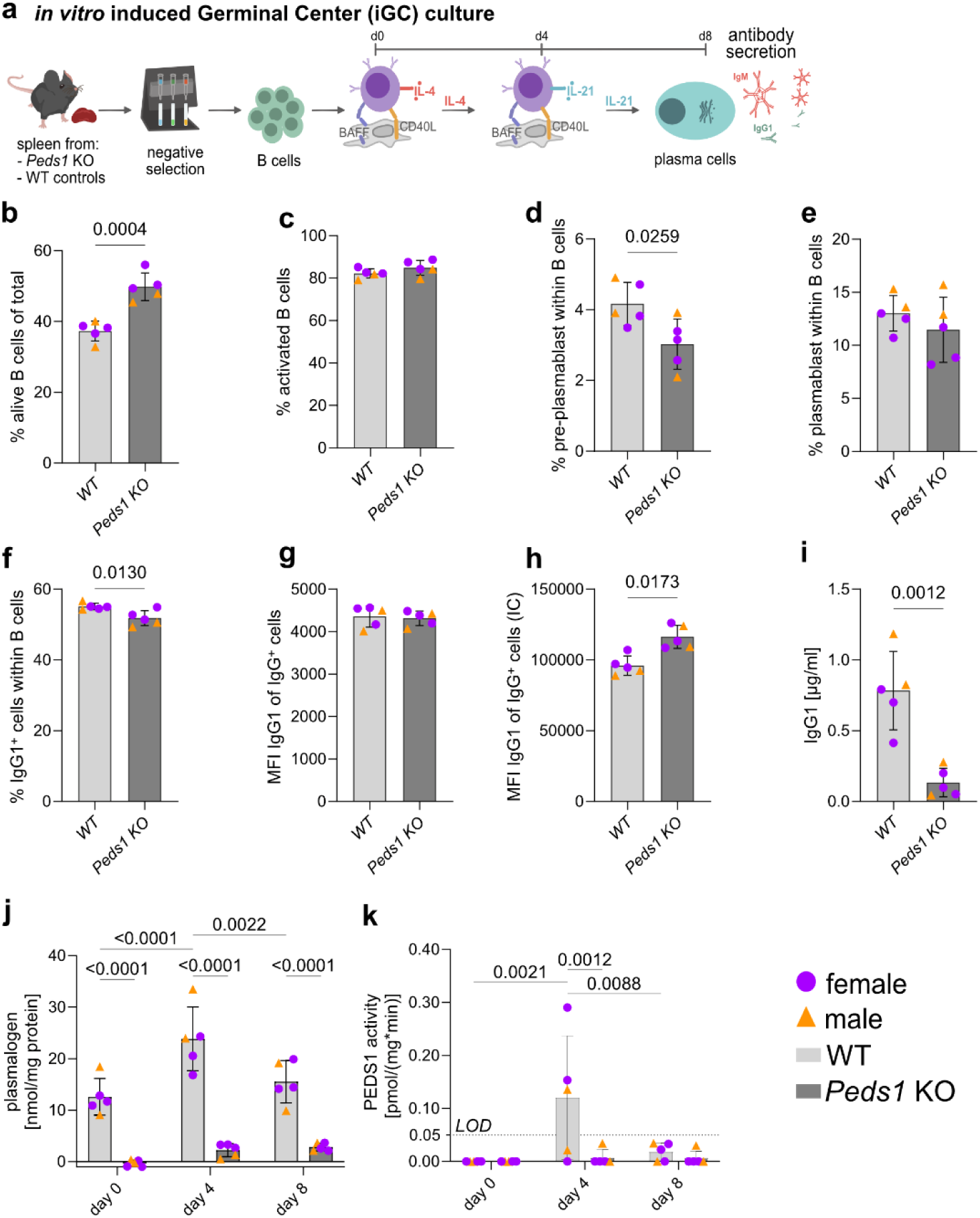
Phenotypic and biochemical profiling of in vitro induced GC B cell cultures derived from splenic B cells of Peds1 KO and WT mice. **a** Workflow of the iGC culture system. **b** Total viable B cell yield at day 8 of culture. **c** Frequency of activated B cells (B220^+^, GL7^+^, CD95^+^) within the total B cell population. **d** Frequency of pre-plasmablasts (B220^+^, CD138-) and **e** plasmablasts (B220^+^, CD138^+^) within the B cell population. **f** Frequency of surface IgG1^+^ cells and **g** mean fluorescence intensity (MFI) of surface IgG1 within the total iGC B cell population. **h** MFI of intracellular IgG1 within the total iGC B cell population. **i** IgG1 quantity in culture supernatant, measured by ELISA. **j** Plasmalogen content and **k** PEDS1 enzymatic activity in FACS-sorted naïve B cells ex vivo and iGC culture-derived B cells on days 4 and 8, measured via HPLC. Females are shown as violet circles and males as yellow triangles. Data are presented as mean ± SD, for both WT and Peds1 KO (n = 5, 3 female and 2 male mice). Statistical comparisons were made using two-tailed Student’s t-test (**b-i**) or two-way ANOVA with Bonferroni’s multiple comparison test (**j-k**).

Despite similar starting B cell numbers, *Peds1* deficiency resulted in a higher total B cell yield on day 8 of the iGC culture (*Peds1* KO = 49.8 ± 3.91, WT = 37.3 ± 2.76, *p* = 0.0004, Fig. 4b), while the overall cell cycle distribution remained unchanged (Suppl. Fig. 5a). On this specific day all B cells could be assigned to one of three phenotypic states: activated B cells, pre-plasmablasts or plasmablasts. The frequency of activated B cells was unchanged (Fig. 4c). In contrast, IL-21-driven progression into the pre-plasmablast state was significantly diminished in *Peds1* KO cultures (*Peds1* KO = 3.03 ± 0.709, WT = 4.17 ± 0.611, *p* = 0.0259, Fig. 4d), while plasmablast frequencies showed only a trend towards reduction (Fig. 4e). Overall, CSR from IgM to IgG1 was modestly reduced, reflected by fewer IgG1^+^ cells in *Peds1* KO cultures (*Peds1* KO = 51.8 ± 2.12, WT = 55.1 ± 0.944, *p* = 0.013, Fig. 4f). Additional flow cytometric analyses revealed unaffected IgE^+^ cell frequency (Suppl. Fig. 5b), and a trend towards an increased IgM^+^ cell fraction with unaltered staining intensity for cell surface or intracellular IgM (Suppl. Fig. 5c-g). IgG1 cell surface staining intensity on IgG1^+^ cells was unchanged (Fig. 4g), however, we found a marked intracellular accumulation of IgG1 (*Peds1* KO = 116,379 ± 8,234, WT = 96,073 ± 6,829, *p* = 0.0173, Fig. 4h). At the same time, IgG1 secretion in the culture supernatant was strongly reduced (Fig. 4i). This aligns with the lower basal IgG1 titres observed at steady state *in vivo* and is compatible with decreased IgG1 secretion by *Peds1* KO culture plasmablasts and/or plasma cells.

Next, we measured plasmalogen levels and PEDS1 enzymatic activity in FACS-sorted total and iGC culture-derived days 4 and 8 B cells, using two established reversed-phase HPLC assays^39^. As expected, *Peds1* KO B cells generally showed a marked reduction in plasmalogens and no PEDS1 activity (Fig. 4j, k). In WT cultures, plasmalogen content during iGC differentiation increased prominently on day 4 (Fig. 4j). Consistent with this, PEDS1 activity peaked at day 4 (Fig. 4k). Together, these data indicate that iGC differentiation is accompanied by a transient rise in plasmalogens that depends on PEDS1 activity.

Collectively, these results demonstrate that *Peds1*-deficient B cells can proliferate and differentiate into plasmablasts, while the secretion of IgG1 is impaired.

### *Peds1* deficiency has an effect on GC dynamics, antibody production, and oxidative stress *in vivo*

Given the phenotypes observed at steady state *in vivo* and *in vitro,* we next examined the consequences of plasmalogen deficiency during an *in vivo* immune challenge. Mice were immunised with ovalbumin (OVA) and peripheral blood was collected at days 0 (pre-immunisation), 7 and 14 for immunoglobulin analysis. Animals were sacrificed on day 14 for haematological, cellular and immunological analyses of spleen and bone marrow (Fig. 5a).

**Figure 5:**
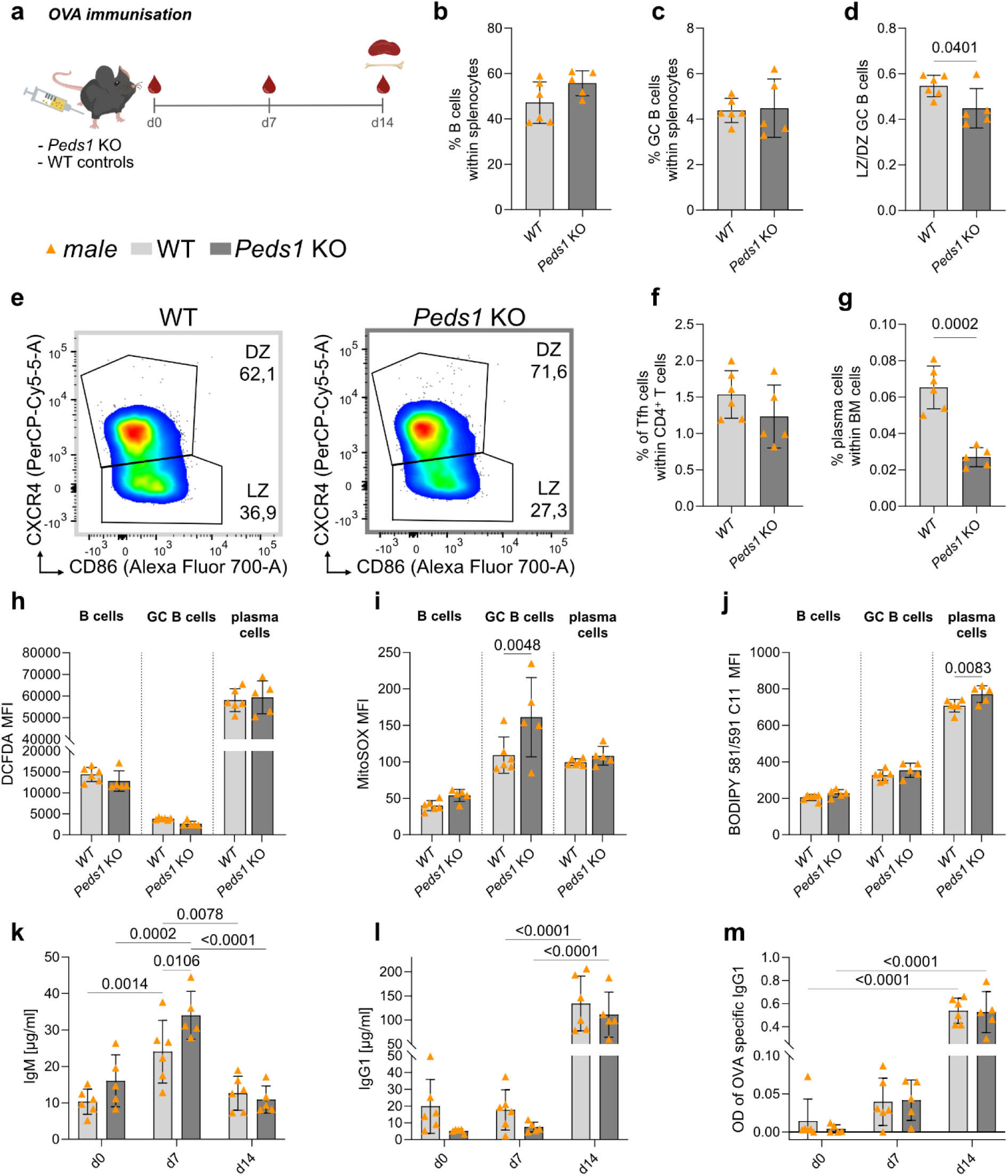
In vivo analysis of GC responses after OVA immunisation of Peds1 KO and WT mice. **a** Scheme of the OVA immunisation and sampling schedule. Blood was collected at day 0, 7 and 14, with terminal harvest of immune organs on day 14. Frequencies of **b** total B cells (B220^+^, CD19^+^) and **c** GC B cells (B220^+^, CD19^+^, CD38^-^) among splenocytes, assessed by flow cytometry. **d** Ratio of LZ to DZ GC B cells (LZ: CXCR4^low^, CD86^high^, DZ: CXCR4^high^, CD86^low^) and **e** representative flow cytometry plots illustrating the gating strategy for LZ and DZ cells of WT and Peds1 KO mice. **f** Frequency of Tfh cells among splenic CD4^+^ T cells (CXCR5^+^, PD-1^+^). **g** Plasma cell frequency (TACI^+^, CD138^+^) in bone marrow. **h** Total cellular ROS (DCFDA) MFI in B cells, GC B cells and plasma cells. **i** Mitochondrial superoxide (MitoSOX) MFI in total B cells, GC B cells and plasma cells. **j** Lipid ROS (BODIPY 581/591 C11) MFI in B cells, GC B cells and plasma cells. Serum **k** IgM and **l** IgG1 levels at days 0, 7, and 14 post-immunisation, quantified by ELISA. **m** OVA-specific IgG1 in serum, reported as optical density (OD). Each data point represents an individual mouse, WT (n = 6 male mice), Peds1 KO (n = 5 male mice). Data are shown as mean ± SD. Statistical comparisons were made using two-tailed Student’s t-test (**b-d, f-g**) or two-way ANOVA with Bonferroni’s multiple comparisons test (**h-m**).

Most readouts were comparable between genotypes after immunisation. We detected no genotype-dependent differences in body weight, spleen/body weight ratio, spleen and bone marrow cell counts (Suppl. Fig. 6a-d).

*Peds1* KO mice displayed a modest, non-significant increase in splenic B cell counts (*Peds1* KO 55.8 ± 5.5, WT 47.22 ± 9.16, *p* = 0.102, Fig. 5b), while they exhibited the expected immunisation-dependent expansion of GC B cells. Thus, GC B cell frequencies remained comparable between genotypes (Fig. 5c). This differed from the steady state setting, where GC B cell frequencies were reduced in mesenteric lymph nodes and Peyer’s patches of *Peds1* KO mice (Fig. 3e, f).

GC B cells cycle between the dark zone (DZ), where they undergo clonal expansion and somatic hypermutation, and the light zone (LZ), where selection occurs in collaboration with Tfh cells^37^. Despite similar overall GC B cell frequencies, *Peds1* KO mice showed a reduced LZ to DZ ratio (Fig. 5d, e), elicited by a relative increase in DZ GC B cells and decrease in LZ GC B cells (Suppl. Fig. 6e and f, respectively). Tfh cell frequency was slightly reduced, similar to the reduction observed in the Peyer’s patches of unchallenged animals (*Peds1* KO 1.23 ± 0.433, WT 1.54 ± 0.328, *p* = 0.219, Fig. 5f), as well as for CD4^+^ T cells (*Peds1* KO = 13.2 ± 1.59, WT = 14.6 ± 1.65, *p* = 0.1908, Suppl. Fig. 6g). Whereas the splenic plasma cell fraction was comparable between genotypes (Suppl. Fig. 6h), bone marrow plasma cell frequencies were significantly reduced in *Peds1* KO mice (*Peds1* KO = 0.027 ± 0.0052, WT = 0.0653 ± 0.0118, *p* = 0.0002; Fig. 5g). Together, these data point to a reduced plasma cell output after immunisation in *Peds1* KO mice.

Building on recent work by Cho *et al.* demonstrating that *Dhrs7b* deficiency, which disrupts ether lipid biosynthesis early in the pathway, increases oxidative stress and alters GC B cell fate decisions and antibody responses^28^, we next assessed oxidative stress parameters in B cells, GC B cells and plasma cells after OVA immunisation. Flow cytometric analysis displayed no genotype-dependent changes in total cellular ROS in B cells, GC B cells or plasma cells (Fig. 5h). In contrast, mitochondrial superoxide was specifically increased in *Peds1* KO GC B cells (*Peds1* KO = 161.0 ± 54.4, WT = 109.0 ± 24.9, *p* = 0.004, Fig. 5i), while this was unvaried in total B or plasma cells. Finally, lipid peroxidation was specifically increased in *Peds1* KO plasma cells (*Peds1* KO = 771.0 ± 46.0, WT = 708.0 ± 34.2, *p* = 0.008, Fig. 5j), while comparable between genotypes in total B and GC B cells. Our results collectively indicate that selective plasmalogen deficiency led to specific changes only in mitochondrial superoxide and oxidised lipid species which did, however, not translate into detectable changes in total cellular ROS.

To test whether the reduced basal IgG1 observed at steady state and the lower IgG1 secretion in the iGC culture translated into an altered Ig response in *Peds1* KO mice upon immunisation *in vivo*, we performed ELISA for serum immunoglobulin levels at day 0 and days 7, and 14 post-immunisation. Total IgM was significantly increased in WT and *Peds1* KO mice at day 7 and returned to baseline levels at day 14. Further, day 7 IgM levels were significantly higher in *Peds1* KO than in WT (Fig. 5k). OVA-specific IgM titres remained unchanged (Suppl. Fig. 6i). The IgG1 levels exhibited a trend toward reduction in *Peds1* KO at days 0 and 7, mirroring the reductions observed in steady state serum and *in vitro* iGC cultures, but converged with WT levels by day 14 (Fig. 5l), when both genotypes showed a significant proficient class switching. OVA-specific IgG1 titres showed a significant increment by day 14 comparably between genotypes (Fig. 5m). Thus, despite lower basal IgG1, *Peds1* KO mice mounted a robust IgG1 response to OVA by day 14.

While still permitting a robust antibody response upon immunisation, our data identify plasmalogen deficiency as a modulator of humoral immunity that shapes GC dynamics and IgG1 output under certain conditions.

## Discussion

The contribution of ether lipid metabolism to immune regulation is only beginning to emerge, and the specific role of plasmalogens in immunity remains essentially unexplored. Serum plasmalogens are reduced in multiple sclerosis and in chronic inflammatory demyelinating polyneuropathy (CIDP) patients^40,41^, which may reflect inflammatory lipid remodelling and/or oxidative consumption and is consistent with a role in oxidative defence. Recently identified recessive *PEDS1* variants in two patients cause neurodevelopmental disorder with recurrent respiratory infections, raising the possibility of immune vulnerability in the setting of selective plasmalogen deficiency^14^. The identification of the gene encoding PEDS1 enables the genetic dissection of ether lipid subclasses and plasmalogen function *in vivo^7^*. Our study identifies plasmalogens and their biosynthetic enzyme PEDS1 as a critical lipid-metabolic modulator that influences GC dynamics, class switching and antibody secretion, while leaving most aspects of haematopoiesis and lymphocyte development largely intact.

In *Peds1*-deficient mice we observed reduced red blood cell counts and haematocrit, as well as increased mean platelet volume. These findings are consistent with previous reports in the same mouse model by the IMPC^35^. We expanded our investigations to bone marrow erythroid differentiation in bone marrow and found no differences in progenitor frequencies. Further, the cellularity of immune organs such as bone marrow, spleen and thymus was not affected by PEDS1 loss. The myeloid compartment was also largely preserved, with mild changes confined to peripheral blood granulocytes, neutrophils, and eosinophils. Together, these data suggest that baseline haematopoietic homeostasis tolerates plasmalogen loss with only limited changes at steady state. On the B cell side, canonical bone marrow and peripheral maturation steps were preserved, yet GC B cells were reduced in gut-associated lymphoid tissues, including mesenteric lymph nodes and Peyer’s patches, and basal IgG1 serum titres were selectively lowered. A recent report by Cho *et al*. implicated loss of all ether lipid subclasses, elicited by knockout of the enzyme DHRS7B and consequent broad ether lipid depletion, in reduced antigen-induced B cell proliferation and GC B cell number^28^. Together with the work from Cho *et al.*, our findings support a model in which plasmalogens are dispensable for lineage commitment, but required for optimal GC responses and IgG1 secretion in steady state.

To further dissect the impact of plasmalogens on the GC, the iGC culture system allowed us to study the requirement of this ether lipid subclass for B cell activation, class switch recombination, plasmablast generation and antibody production *in vitro*. Switching to IgG1 was modestly reduced, and intracellular IgG1 accumulation coincided with hampered IgG1 release into the supernatant. The dynamic rise in plasmalogen content and PEDS1 activity in WT iGC cultures, which was absent in *Peds1* KO cultures, supports the idea that plasmalogen biosynthesis is engaged during iGC differentiation. This cannot be readily compensated by the accumulation of plasmanyl lipids, which are the precursors for plasmalogen synthesis and build up upon *Peds1* deficiency^28,34^. The preservation of plasmablast generation alongside reduced IgG1 release suggests a role for plasmalogens in membrane organisation, vesicular trafficking, or secretory pathway fitness. Such effects would be expected to preferentially impact high-rate antibody secretion, rather than early activation or proliferation *per se*. In a different cell system, Mansell *et al.*, reported that AGPS KO-driven ether lipid deficiency in pB3 murine breast carcinoma cells alters membrane properties, including fluidity and tension^42^.

Our *in vivo* OVA immunisation experiments further refine this picture, by revealing how plasmalogen loss is associated with changes in GC zoning and redox balance under antigen-specific immune challenge. Despite unaltered overall splenic GC frequencies, *Peds1* deficiency shifted GC zoning toward the dark zone. Concurrently, bone marrow plasma cell numbers were diminished, indicating compromised humoral output. This phenotype was also found by Cho *et al.*, who reported that total ether lipids deficiency via B cell-specific DHRS7B deletion, not only reduced GC size and numbers after immunisation, but also impaired the magnitude and affinity maturation of NP-specific IgG1 responses in the NP-OVA immunisation model^28^. Unlike upstream ether lipid blockade, which curtails GC formation and expansion, plasmalogen loss maintained GC frequencies but altered the LZ–DZ equilibrium, shifting GC B cells towards the DZ. This shift may reflect altered cycling between DZ expansion and LZ selection, with potential consequences for downstream plasma cell output. Collectively, the data delineate that plasmalogens modulate GC organisation and downstream plasma cell output during immune challenge, in a manner that is separable from broader ether lipid requirements for GC expansion.

We also conducted ROS studies after OVA immunisation of *Peds1* KO mice compared to WT controls to evaluate the impact of selective plasmalogen loss on cellular redox homeostasis. Total cellular ROS levels were unchanged in B cells, GC B cells and plasma cells, whereas mitochondrial superoxide increased in GC B cells and lipid peroxidation was elevated specifically in plasma cells. This compartment-specific pattern is consistent with distinct redox pressures across the GC-to-plasma cell axis under plasmalogen deficiency. Given that plasmalogens can act as membrane-associated redox buffers via their vinyl ether double bond^5,43^ and can influence membrane organisation^15,40,43^, their loss may sensitise GC B cells and plasma cells to oxidative stress during intense proliferation and antibody synthesis, while leaving bulk cytosolic ROS levels largely unchanged^28,44^.

We also found that antigen-specific IgG1 responses to OVA reached WT-like titres at day 14 after immunisation in *Peds1* KO mice, indicating that, in contrast to the *in vitro* setting, compensatory mechanisms present *in vivo* can partially restore humoral performance, despite altered GC zoning and redox homeostasis. The *in vivo* compensation in our *Peds1*-deficient mice after OVA immunisation contrasts with the observations of Cho *et al.*, who observed persistent reduction in serum NP-specific antibody production, including IgG1, even after boosting^28^.

Taken together, our findings position plasmalogens and PEDS1 for the first time as context-dependent modulators of B cell immunity. Plasmalogens appear to be largely dispensable for baseline haematopoiesis and lymphocyte development, but contribute to optimal GC organisation, IgG1 class switching, and efficient antibody secretion under conditions of high metabolic and oxidative demand. The selective nature of the defects, restricted to particular antibody isotypes, GC polarisation, and plasma cell redox control, suggests that ether lipid pathways may be exploitable to modulate humoral immunity, without broadly compromising haematopoietic integrity.

The role of ether lipids in membrane fusion^45,46^, as well as in membrane plasticity and fluidity^42^, and their restoration via supplementation in deficiency models^5^ further supports the idea that ether lipids can influence secretory dynamics and may have therapeutic relevance. Future work should clarify how plasmalogen-driven membrane and organelle remodelling intersect with B cell receptor signalling, secretory pathway dynamics, and redox-sensitive cell death pathways in GC B cells and plasma cells.

## Methods

### Ethics statement and animal models

All animal breeding colonies and experimental protocols received official approval from the Austrian Federal Ministry of Education, Science and Research (GZ 2024-0.307.678 and GZ 2025-0.247.787). We acquired the *Peds1* KO mouse model (strain designation: Tmem189tm1a(KOMP)Wtsi) from the Wellcome Sanger Institute (Hinxton, Cambridge, UK) distributed through the European Mouse Mutant Archive EMMA/Infrafrontier GmbH (Munich, Germany)^47–50^. The mice were bred and maintained on a C57bl/6N genetic background. This transgenic line features a knockout-first allele designed to produce an inactive, truncated PEDS1 protein. Mouse genotyping and genotype verification of the experimental animals was performed as described previously^34^. Mice were maintained under specific pathogen free (SPF) conditions at the animal facility of the Medical University of Innsbruck and were housed in individually ventilated cages equipped with nesting material. The animals were kept under a controlled 12 h/12 h light/dark cycle, with 21°C ambient temperature and 50% humidity, and were provided with *ad libitum* standard chow and water. Both female and male littermate mice aged 8 weeks (82-85 days) were analysed, and sex was considered in the analysis of the individual parameters. The immunisation experiment was conducted only on male mice aged 7-9 weeks and consequently their age at the endpoint of the study was 9-11 weeks.

### Single-cell suspension preparation

For the preparation of single-cell suspensions and the subsequent washing steps, a staining buffer containing PBS, 2% FBS (Thermo Fisher Scientific, Paisley, United Kingdom, 10270-106) and 10 µg/ml gentamicin (Thermo Fisher Scientific, 15750037), was used. Murine spleen, mesenteric lymph nodes and Peyer’s patches were mechanically dissociated by meshing them through a 70 µm cell strainer (Merck, Darmstadt, Germany, 352350), whereas bone marrow was collected from femur and tibia by flushing the bones with a 23G needle. For the peritoneal lavage, the peritoneal cavity was injected *post mortem* with 10 ml of staining buffer using a 27G needle and, after a gentle massage, the fluid was extracted with the same needle.

Peripheral blood was collected from the *vena submandibularis* into tubes coated with 8 µl heparin (Gilvasan Pharma GmbH, Vienna, Austria, 5000 i.E./ml). Blood analysis was performed using a blood cell counter (scil animal care company GmbH, Viernheim, Germany). For flow cytometry analysis of spleen, red blood cells were depleted by resuspending cell pellets in 1 ml of lysis buffer (155 mM NH_4_Cl, 10 mM KHCO_3_, 0.1 mM EDTA; pH 7.5), followed by incubation on ice for 3 min. Two consecutive 5 min incubation steps were performed. The cells were then washed with the staining buffer and filtered through a 50 µm cup falcon (BD Biosciences, San Diego, CA, USA, 340632). Cell counts and viability were determined using a counting chamber (Merck, BR717805) and trypan blue exclusion.

### Flow cytometry analysis

Cells were analysed using a BD Bioscience LSRII-Fortessa flow cytometer. The fluorochrome-labelled antibodies which were used are listed in Table 1. Quantitative data analysis was performed using FlowJo (FlowJo v10; BD, Ashland, OR, USA), which included doublets exclusion. A minimum of 100,000 cells per sample was acquired. Biotin-conjugated antibodies were incubated with fluorochrome-labelled streptavidin conjugates at a dilution of 1:200. For the cell death analysis, cells were stained immediately before measurement by adding Annexin V-FITC (BioLegend, San Diego, CA, USA, 640945, 1:1,000) or TO-PRO™-3 Iodide 642/661 (1:50,000, Thermo Fisher Scientific, T3605). These stains were diluted in Annexin V Binding Buffer (eBioscience, San Diego, CA, USA, 00-0055-56, 1:10 dilution in water).

**Table 1:** List of all fluorescent-labelled anti-mouse antibodies used for flow cytometry.

| Channel | Antibody | Provider | Cat. Number | Clone | Dilution |
| --- | --- | --- | --- | --- | --- |
| **Alexa Fluor 700** | α CD21/CD35 (CR2/CR1) | BioLegend | 123431 | 7 E 9 | 200 |
|  | α CD3ε | BioLegend | 152316 | 500A2 | 200 |
|  | α I-A/I-E | BioLegend | 107622 | M5/114.15.2 | 200 |
|  | α IgG1 | BioLegend | 406632 | RMG1-1 | 100 |
| **APC** | Streptavidin | BioLegend | 405207 | - | 200 |
|  | α CD11c | BioLegend | 117309 | N418 | 200 |
|  | α CD71 (Transferrin receptor) | Invitrogen | 17-0711-80 | R17217 | 200 |
|  | α CD267 (TACI) | BD | 558453 | 8F10 | 100 |
|  | α CD93 (AA4.1, early B lineage) | BioLegend | 136510 | AA4.1 | 200 |
| **APC/Cyanine7** | α CD23 | BioLegend | 101629 | B3B4 | 100 |
|  | α CD4 | BioLegend | 100414 | GK1.5 | 200 |
|  | α Ly-6A/E (Sca-1) | BioLegend | 108126 | D7 | 200 |
|  | α CD11b | BioLegend | 101226 | M1/70 | 200 |
| **Biotin** | α CD11b | Miltenyi Biotec | 130-113-242 | REA592 | 200 |
|  | α CD185 (CXCR5) | BioLegend | 145510 | L138D7 | 100 |
|  | α CD19 | BioLegend | 115504 | 6D5 | 200 |
|  | α F4/80 | BioLegend | 123105 | BM8 | 200 |
|  | α NK-1.1 | BioLegend | 108704 | PK136 | 200 |
|  | α CD45R/B220 | BioLegend | 103204 | RA3-6B2 | 200 |
| **Brilliant Violet 421** | α CD117 (c-kit) | BioLegend | 105828 | 2B8 | 300 |
|  | α CD5 | BioLegend | 100617 | 53-7.3 | 100 |
|  | α CD8a | BioLegend | 100738 | 53-6.7 | 200 |
|  | α Ly-6G | BioLegend | 127627 | 1A8 | 200 |
| **Brilliant Violet 510** | α CD138 (Syndecan-1) | BioLegend | 142521 | 281-2 | 200 |
|  | α CD44 | BioLegend | 103044 | IM7 | 200 |
| **Brilliant Violet 605** | Streptavidin | BioLegend | 405229 | - | 200 |
|  | α CD150 (SLAM) | BioLegend | 115927 | TC15-12F12.2 | 200 |
|  | α CD19 | BioLegend | 115540 | 6D5 | 200 |
| **Brilliant Violet 650** | α CD19 | BioLegend | 115555 | 6D5 | 200 |
|  | α CD11b | BioLegend | 101259 | M1/70 | 200 |
|  | α CD45R/B220 | BioLegend | 103246 | RA3-6B2 | 100 |
| **eFluor 450** | α IgM | eBioscience | 48-5890-82 | eB121-15F9 | 100 |
| **FITC** | α CD19 | BioLegend | 115506 | 6D5 | 200 |
|  | α CD38 | BioLegend | 102705 | 90 | 200 |
|  | α FcεRIα | BioLegend | 134305 | MAR-1 | 200 |
|  | α TCR β chain | BioLegend | 109206 | H57-597 | 100 |
| **PE** | Streptavidin | eBioscience | 12-4317-87 | - | 200 |
|  | α CD170 (Siglec-F) | BioLegend | 155505 | S17007L | 300 |
|  | α CD1d | eBioscience | 12-0011-83 | 1B1 | 100 |
|  | α CD279 (PD-1) | BD Biosciences | 551892 | J43 | 300 |
|  | α CD43 | BD Biosciences | 553271 | S7 | 200 |
| **PE/Cyanine7** | α CD115 (CSF-1R) | BioLegend | 135523 | AFS98 | 300 |
|  | α CD23 | BioLegend | 101613 | B3B4 | 200 |
|  | α CD48 | BioLegend | 103423 | HM48-1 | 100 |
|  | α IgM | BioLegend | 406514 | RMM-1 | 200 |
|  | α NK-1.1 | BioLegend | 108713 | PK136 | 200 |
| **PerCP/Cyanine5.5** | α CD62L | BioLegend | 104431 | MEL-14 | 300 |
|  | α IgD | BioLegend | 405710 | 11-26.2a | 200 |
|  | α Ly-6C | BioLegend | 128012 | HK1.4 | 200 |
|  | α TER-119/Erythroid Cells | BioLegend | 116228 | TER-119 | 200 |

### Enzyme-linked immunosorbent assay (ELISA)

To quantify Igs in mouse serum and iGC culture supernatant ELISA was performed. For serum preparation, blood was collected using lancets (Thermo Fisher Scientific, NC9891620) from the submandibular vein and kept in tubes at 4°C for at least 2 h, and centrifuged at 12,000 g for 5 min at 4°C. The supernatant was stored at -80°C until further use. 96-well ELISA plates (Merck KGaA, Darmstadt, Germany, CLS3590) were coated with 100 µl of 50 µg/ml capture antibody (Southern Biotech, Birmingham, AL, USA, 1010-01, 50 μg/ml in PBS) at 4°C overnight. For ovalbumin (OVA)-specific analysis of IgM and IgG1 after immunisation, the plates were coated with 100 µl of 10 μg/ml OVA (Merck, A5503). Post-coating, the plates were washed three times, blocked for 4 h at RT with blocking buffer (1x PBS with 1% BSA) and washed again. The coated wells were incubated at 4°C overnight with either mouse serum (diluted 1:25-1:2,000 for total IgM, IgG1 and IgG2c detection, 1:100 for OVA-specific IgM and IgG1) or iGC *in vitro* culture supernatant (1:100 dilution). Following incubation, the plates were washed three times with washing buffer (1x PBS with 0.05% Tween20). HRP-conjugated anti-mouse IgG1, IgG2c or IgM (Southern Biotech, Birmingham, AL, USA, 1070-05 and 1020-05, 1:5,000 in ELISA blocking buffer) was added to the wells for 4 h at RT. Detection was achieved by adding the ABTS substrate and after a 20-minute incubation at RT, absorbance at 405 nm was measured in a CLARIOStar plate reader (BMG LABTECH, Ortenberg, Germany). Ig concentrations were quantified by interpolation of the measured absorbance values from the standard curve values.

### Induced Germinal Centre (iGC) B cell culture

To study the differentiation of naïve B lymphocytes into plasmablasts *in vitro*, we made use of an induced germinal centre (iGC) co-culture system. This system uses 40LB fibroblast feeder cells, a BALB/3T3 line stably expressing murine CD40L and BAFF^38^ obtained from RIKEN BRC (RIKEN BioResource Research Center, Tsukuba, Ibaraki, Japan). B cells were isolated from the splenic single-cell suspension. This was achieved using MagniSort^TM^ Streptavidine Negative Selection Beads (Thermo Fisher Scientific, MSNB-6002-74) in conjunction with biotinylated antibodies targeting Ter119 (BioLegend, 116204, 1:100), CD11b (BioLegend, 101204, 1:100) and TCRβ (BioLegend, 109204, 1:50), as per manufacturers instructions. The iGC B cell culture followed our adapted protocol^51^. Initially, 4 x 10⁵ 40LB feeder cells were treated with 10 µg/ml mitomycin C (Merck, M0305) in 1 ml of DMEM (Merck, D6429) supplemented with 10% FBS (Gibco, Thermo Fisher Scientific, Waltham, MA, USA, 10270-106), 2 mM L-glutamine (Merck, G7513), and 100 µg/ml penicillin/100 μg/ml streptomycin (Merck, 0781). After treatment, 40LB cells were washed three times with PBS, and subsequently, 7.5 x 10⁵ B cells were added to the feeder cells in B cell medium which consisted of DMEM supplemented with 10% FBS, 2 mM L-glutamine, 10 mM Hepes (LONZA, Basel, Switzerland, BE17-737E), 1 mM sodium pyruvate (Gibco, 13360-039), 1x nonessential amino acids (Gibco, 11140-035), 100 µg/ml penicillin/100 μg/ml streptomycin, 50 μM β-mercaptoethanol (Merck, M3148), and 10 ng/ml recombinant IL-4 (Peprotech, Cranbury, NJ, USA, 214-14). On day 4, the iGC B cells were collected by aspirating the culture medium and washing the plates with harvest buffer (PBS containing 0.5% BSA and 2 mM EDTA). A portion of these cells were either prepared for analysis, and 7.5 x 10⁵ iGC B cells were replated onto new 6-well plates containing fresh mitomycin C-treated 40LB feeder cells. These replated cells were cultured in 4 ml of B cell medium supplemented with 10 ng/ml recombinant IL-21 (Peprotech, 210-21). On day 8, the B cells were harvested for subsequent flow cytometric analysis. For surface flow cytometric analysis, iGC B cells underwent staining according to the general flow cytometry protocol in the section *“Flow cytometry analysis”*. The antibodies used for the staining are listed in Table 1. Prior to data acquisition, cells were labelled with Fixable Viability Dye eFluor 780 (Thermo Fisher Scientific, 65-0865-14), following the manufacturer’s instructions. For intracellular flow cytometric analysis, iGC B cells were first washed once with PBS. Next, they were resuspended in trypsin for 10 minutes and left at 37°C. Cells were washed with staining buffer and fixed by adding a custom-made fixation solution (PBS + 4% PFA + 0.1% saponin) for 20 minutes at 4°C. Subsequently, cells underwent two washes with Perm/Wash buffer (composed of PBS, 1% BSA, 0.1% saponin, and 0.025% sodium azide). Following these washes, cells were incubated for 15 minutes with a primary antibody cocktail. After another wash with Perm/Wash buffer (BD Biosciences, 554723), a secondary antibody solution containing streptavidin BV605 was added, and cells were incubated for an additional 15 minutes. The cells were processed further for flow cytometric analysis, as detailed in the *“Flow cytometry analysis”* section.

### PEDS1 enzymatic activity and plasmalogen content of iGC sorted B cells

For plasmalogen content and PEDS1 enzymatic assay analysis, GC B cells from the *in vitro* iGC culture were harvested by repeated vigorous pipetting. The cell suspension was incubated with anti-CD16/32 (Fc block, 1 µg/ml in 500 µl staining buffer, 10 min), washed, and subsequently labelled for 20 min with antibodies in staining buffer. The cells harvested at day 0 prior to the iGC culture were *ex vivo* naïve cells, whereas the GC B cell population at day 4 and 8 of culture was defined as CD19^+^/B220^+^/CD38^low^/Fas^+^. One million cells from the *in vitro* iGC culture were sorted on a FACSAria III (BD Biosciences) into staining buffer, centrifuged at 541 g for 3 min at 4°C, the supernatant discarded, snap-frozen and stored at -80°C, until further use for the biochemical assays.

PEDS1 enzymatic activity, as well as plasmalogen abundance, was analysed in the sorted iGC B cells following a protocol as described before^39^, with one modification explained below. Briefly, the sorted cell samples were homogenised in 200 µl 0.1 mM Tris HCl and 0.25 M sucrose (pH 7.2), adding 2 micro spoons of glass beads (Merck, G1277), for 4 x 30 s at 20 Hz using a mixer mill (Resch GmbH, Haan, Germany). Protein concentration was quantified by Bradford assay (Bio-Rad, Hercules, California, USA, #5000006) using BSA as standard. The protein concentration of the homogenates ranged from 0.18 to 0.67 mg/ml. In contrast to the published method, both the PEDS1 enzymatic assay and the plasmalogen quantification were performed using the same homogenate, which was not further diluted, instead, final results were normalised to the total protein content.

For PEDS1 enzymatic activity we added 10 µl of a mix containing: 1 M Tris HCl (pH 7.2), 1 mg/ml catalase (Merck, C1345), 8.33 mg/ml NADPH, 100 mM EDTA, 2 μM fluorescent-labelled lyso-substrate (1-*O*-pyrenedecyl-*sn*-glycero-3-phosphoethanolamine) and water directly to 15 µl of the homogenate. This was incubated for 30 min at 37°C and the mixture was subsequently divided into two 10 μl aliquots: one treated with 30 μl of acetonitrile (ACN)/2 M HCl (895:105, v/v) for vinyl ether bond cleavage and the other aliquot (control) was treated with 30 μl of ACN/2 M acetic acid (895:105, v/v), leaving the vinyl ether double bond intact. Both mixtures were incubated for a further 30 min at 37°C. After centrifugation at 20,000 g for 5 min, 10 μl of the supernatant was analysed on an Agilent 1200 HPLC system, equipped with a thermostatted autosampler, UV-Vis and fluorescence detectors and a column thermostat. A Zorbax Eclipse XDB-C8 4.6 × 50 mm, 3.5 μm particle size column (Agilent Technologies, Vienna, Austria) was eluted with 10 mM potassium phosphate buffer (pH 6.0) containing 79% (v/v) methanol for 3 min, at a flow rate of 1 ml/min, followed by a linear gradient to 100% methanol at 10 min. Methanol (100%) was held until 15 min, and the column was then equilibrated to the starting elution buffer until 17 min. The pyrene-labelled compounds were detected by fluorescence (excitation 340 nm, emission 405 nm). Plasmalogen-derived pyrenedecanal liberated upon HCl treatment was monitored by comparison to as external synthetic standard.

To quantify the plasmalogen content of the sorted GC B cells the rest of the homogenate was used for lipid extraction, performed as described before^39^, without a further dilution step. Briefly, to obtain the lipid extracts, the homogenates underwent a double extraction using 500 μl of a chloroform/methanol mixture (2:1, v/v). After combining the organic phases and evaporating them to dryness, the resulting lipid extracts were reconstituted in 100 μl of ACN/ethanol (1:1, v/v). Solubilisation was completed by incubating the samples for 10 min at 37°C with constant agitation. From this, two parallel 10 μl aliquots of the lipid extract were mixed with 40 μl of 0.45 mg/ml dansylhydrazine (Merck, 03334, 0.45 mg/ml), prepared either in ACN/2 M HCl (930:70, v/v), for total aldehyde determination (plasmalogen cleavage) or in ACN/2 M HAc (930:70, v/v), as a control for free aldehydes, which leaves plasmalogens intact. The two parallel samples were incubated for 15 min on ice in the dark, then centrifuged for 5 min at 20,000 g. The resulting supernatants were directly injected to the HPLC as described above and the dansylhydrazine derivatives eluting between 8 and 10 min were quantified by fluorescence peak area (excitation 340 nm, emission 525 nm). Final plasmalogen levels were calculated by correcting the total aldehyde signal for the minor contribution of free aldehydes (typically less than 1%). Octadecanal (Merck, Vienna, Austria) and hexadecanal^52^ were used as external standards for the HPLC analysis after derivatisation with dansylhydrazine.

### *In vivo* ovalbumin immunisation and flow cytometric analysis

For the *in vivo* immunisation, 7-9 weeks old male mice received an intraperitoneal (i.p.) injection of 100 µg Endofit ovalbumin (Invivogen, San Diego, CA, USA, 9006-59-1) mixed in a 1:1 ratio with ImjectTM Alum Adjuvant (Thermo Fisher Scientific, 77161). The total injected volume was 200 µl per mouse. Prior to immunisation at day 0, as well as at days 7 and 14 post-immunisation, blood was collected from the submandibular vein using lancets without analgesia (Thermo Fisher Scientific, NC9891620), for serum Ig analysis via ELISA (see *“Enzyme-linked immunosorbent assay (ELISA)”* methods section). On day 14, mice were sacrificed and organs were harvested for flow cytometry analysis of bone marrow and spleen single-cell suspensions (as detailed above). To assess intracellular ROS, including overall ROS, as well as selective mitochondrial superoxide or lipid peroxide levels in B cells, GC B cells, and plasma cells, splenic single-cell suspensions were prepared after immunisation (see *“Single-cell suspension preparation”* methods section). Cells were first stained with primary and secondary antibodies for cell-type identification (see Table 2). Subsequently, cells were incubated for 30 minutes at 37°C in staining buffer containing either 5 µM of 2’,7’-dichlorodihydrofluorescein diacetate (DCFDA) (for total ROS), 5 µM MitoSOX™ (for mitochondrial superoxide), or 1 µM BODIPY™581/591 C11 (for lipid peroxides). After incubation, cells were washed once with staining buffer and further resuspended in staining buffer containing 10 µg/ml DAPI (Merck, D9542), to allow dead cell exclusion.

**Table 2:**
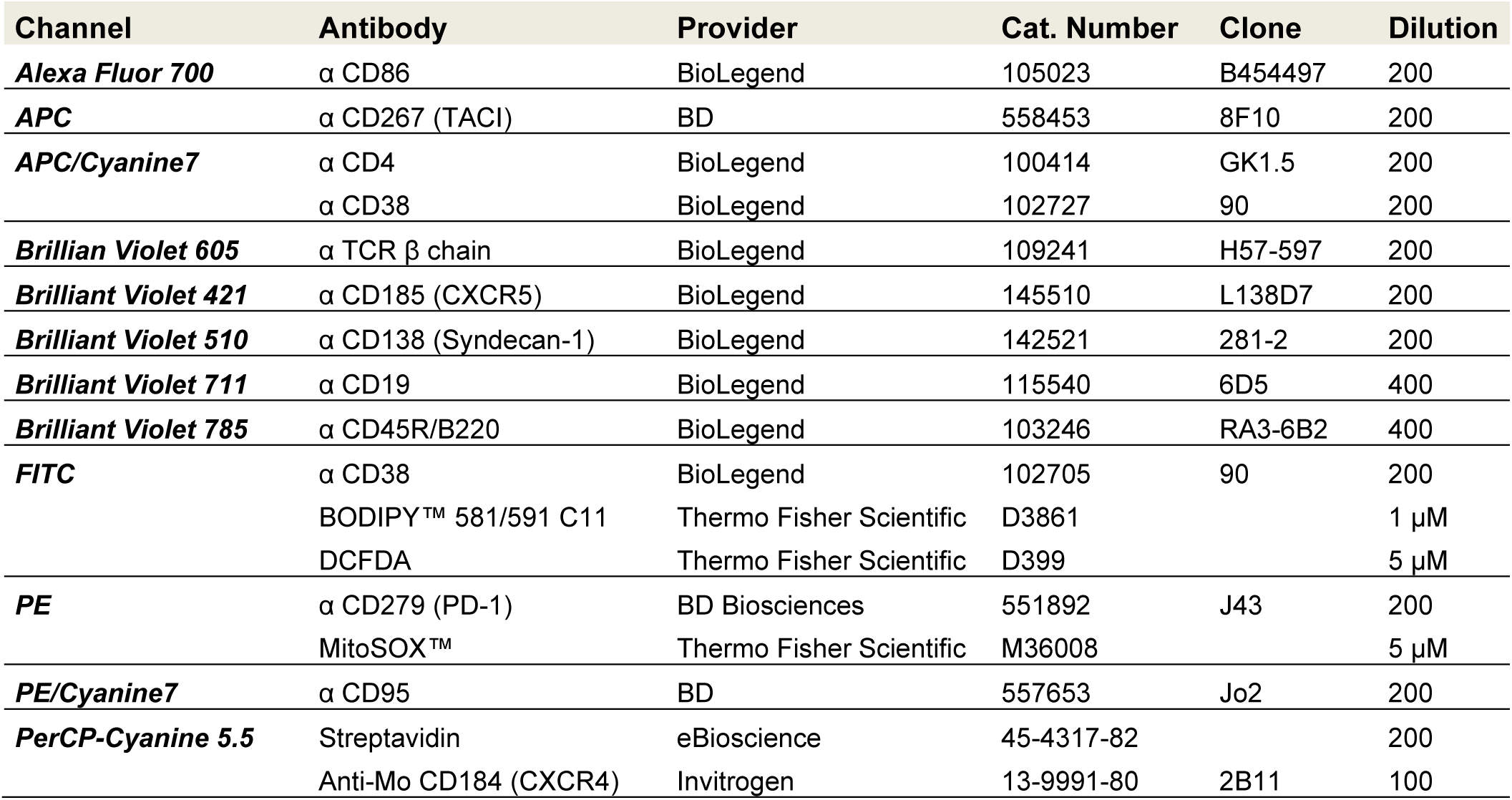
Flow cytometry antibodies used for the immunophenotyping of the in vivo immunisation study.

### Data presentation and statistical analysis

Statistical analyses were conducted using GraphPad Prism version 10.4.1 (GraphPad Software, San Diego, CA, USA). Results are presented as mean ± standard deviation (SD). Comparisons between two groups were assessed using a two-tailed unpaired t-test. For experiments involving multiple groups, one-way ANOVA with Bonferroni post hoc test was applied; for comparisons across multiple conditions or time points, two-way ANOVA followed by Bonferroni multiple comparison correction was used. The number of biological replicates (n) is indicated in each figure legend. Statistical significance was defined as p < 0.05 and exact p values are reported directly in the figures. No statistical methods were used to predetermine sample size. Data collection and analysis were randomised, and investigators were blinded to group allocation during experiments and outcome assessment. Flow cytometry data were processed and analysed using FlowJo v10.9.0 software (BD Biosciences), and all figures were assembled in Affinity Designer 1.10 600bf (Serif Europe Ltd).

## Acknowledgements

The authors thank Nina Madl, Nikolas Andresen, Petra Loitzl, Irene Gaggl, Julia Heppke, Claudia Soratroi, Nadine Heinrich and Martin Saurwein for their expert technical help. We also thank the Wellcome Trust Sanger Institute Mouse Genetics Project (Sanger MPG) and its funders for providing the mutant mouse line Tmem189tm1a(KOMP)Wtsi and INFRAFRONTIER/EMMA (https://www.infrafrontier.eu/). This research was funded in whole or in part by the Austrian Science Fund (FWF) [10.55776/P34723 and 10.55776/FG15 (to K.W.); 10.55776/*P32755* and 10.55776/FG25 to V.L.]. For open access purposes, the author has applied a CC BY public copyright license to any author accepted manuscript version arising from this submission. I.D. acknowledges support by Szabo Scandic (“Immunis Sponsorship for Young Science 2022”).

## Data availability

The datasets generated during this study are available from the corresponding authors upon reasonable request.

## CRediT author Statement

Conceptualisation: I.D, V.L., K.W

Methodology: I.D., S.S., D.K., K.L

Software: I.D., S.S.

Validation: I.D., S.S

Formal analysis: I.D.

Investigation: I.D., S.S., D.K., K.L., F.E., M.A.S.

Resources: V.L.

Data curation: I.D., S.S., V.L., K.W.

Writing - original draft: I.D., K.W.

Writing - Review & Editing: I.D, S.S., D.K., K.L., F.E., M.A.S., G.G., V.L., K.W.

Visualisation: I.D.

Supervision: G.G., V.L., K.W.

Project administration: V.L., K.W.

Funding acquisition: I.D., V.L., K.W.

## References

1. Liebisch, G. et al. Update on LIPID MAPS classification, nomenclature, and shorthand notation for MS-derived lipid structures. J. Lipid Res. 61, 1539–1555 (2020).

2. Sud, M. et al. LMSD: LIPID MAPS structure database. Nucleic Acids Res. 35, D527–32 (2007).

3. Koch, J., Watschinger, K., Werner, E. R. & Keller, M. A. Tricky isomers-the evolution of analytical strategies to characterize plasmalogens and plasmanyl ether lipids. Front. Cell Dev. Biol. 10, 864716 (2022).

4. Koch, J. et al. Benchmarking of trapped Ion Mobility Spectrometry in differentiating plasmalogens from other ether lipids in lipidomics experiments. Anal. Chem. 97, 10578–10587 (2025).

5. Dorninger, F., Werner, E. R., Berger, J. & Watschinger, K. Regulation of plasmalogen metabolism and traffic in mammals: The fog begins to lift. Front Cell Dev Biol 10, 946393 (2022).

6. Gallego-García, A. et al. A bacterial light response reveals an orphan desaturase for human plasmalogen synthesis. Science 366, 128–132 (2019).

7. Werner, E. R. et al. The *TMEM189* gene encodes plasmanylethanolamine desaturase which introduces the characteristic vinyl ether double bond into plasmalogens. Proc. Natl. Acad. Sci. U. S. A. 117, 7792–7798 (2020).

8. Wainberg, M. et al. A genome-wide atlas of co-essential modules assigns function to uncharacterized genes. Nat Genet 53, 638–649 (2021).

9. Steinberg, S. J. et al. Peroxisome biogenesis disorders. Biochim. Biophys. Acta 1763, 1733–1748 (2006).

10. Braverman, N., Argyriou, C. & Moser, A. Human disorders of peroxisome biogenesis: Zellweger spectrum and rhizomelic chondrodysplasia punctata. in Molecular Machines Involved in Peroxisome Biogenesis and Maintenance 63–90 (Springer Vienna, Vienna, 2014).

11. Duker, A. L. et al. Rhizomelic chondrodysplasia punctata morbidity and mortality, an update. Am. J. Med. Genet. A 182, 579–583 (2020).

12. Wanders, R. J. et al. Age-related differences in plasmalogen content of erythrocytes from patients with the cerebro-hepato-renal (Zellweger) syndrome: implications for postnatal detection of the disease. J. Inherit. Metab. Dis. 9, 335–342 (1986).

13. Fazi, C., et al. Case report: Zellweger syndrome and humoral immunodeficiency: The relevance of newborn screening for primary immunodeficiency. Front. Pediatr. 10, 852943 (2022).

14. Amelan, A. et al. CRISPR knockout screens reveal genes and pathways essential for neuronal differentiation and implicate PEDS1 in neurodevelopment. Nat. Neurosci. 1–12 (2026).

15. Braverman, N. E. & Moser, A. B. Functions of plasmalogen lipids in health and disease. Biochim. Biophys. Acta 1822, 1442–1452 (2012).

16. Iannitti, T. & Palmieri, B. An update on the therapeutic role of alkylglycerols. Mar. Drugs 8, 2267–2300 (2010).

17. Arroyo, A. B. et al. Peds1 deficiency in zebrafish results in myeloid cell apoptosis and exacerbated inflammation. Cell Death Discov. 10, 388 (2024).

18. Rubio, J. M., Astudillo, A. M., Casas, J., Balboa, M. A. & Balsinde, J. Regulation of phagocytosis in macrophages by membrane ethanolamine plasmalogens. Front. Immunol. 9, 1723 (2018).

19. Jalil, A. et al. Plasmalogen remodeling modulates macrophage response to cytotoxic oxysterols and atherosclerotic plaque vulnerability. Cell Rep. Med. 6, 102131 (2025).

20. Wallner, S. et al. Phosphatidylcholine and phosphatidylethanolamine plasmalogens in lipid loaded human macrophages. PLoS One 13, e0205706 (2018).

21. Lodhi, I. J. et al. Peroxisomal lipid synthesis regulates inflammation by sustaining neutrophil membrane phospholipid composition and viability. Cell Metab. 21, 51–64 (2015).

22. Lodhi, I. J., Link, D. C. & Semenkovich, C. F. Acute ether lipid deficiency affects neutrophil biology in mice. Cell Metab. 21, 652–653 (2015).

23. Dorninger, F., Wiesinger, C., Braverman, N. E., Forss-Petter, S. & Berger, J. Ether lipid deficiency does not cause neutropenia or leukopenia in mice and men. Cell Metab. 21, 650–651 (2015).

24. Triggiani, M., Schleimer, R. P., Warner, J. A. & Chilton, F. H. Differential synthesis of 1-acyl-2-acetyl-sn-glycero-3-phosphocholine and platelet-activating factor by human inflammatory cells. J. Immunol. 147, 660–666 (1991).

25. Dorninger, F., Forss-Petter, S., Wimmer, I. & Berger, J. Plasmalogens, platelet-activating factor and beyond - Ether lipids in signaling and neurodegeneration. Neurobiol. Dis. 145, 105061 (2020).

26. O’Donnell, V. B., Rossjohn, J. & Wakelam, M. J. Phospholipid signaling in innate immune cells. J. Clin. Invest. 128, 2670–2679 (2018).

27. Balsinde, J. & Balboa, M. A. Plasmalogens in innate immune cells: From arachidonate signaling to ferroptosis. Biomolecules 14, 1461 (2024).

28. Cho, S. H. et al. B cell expression of the enzyme PexRAP, an intermediary in ether lipid biosynthesis, promotes antibody responses and germinal center size. bioRxivorg 2024.10.17.618760 (2024).

29. Facciotti, F. et al. Peroxisome-derived lipids are self antigens that stimulate invariant natural killer T cells in the thymus. Nat. Immunol. 13, 474–480 (2012).

30. Wajda, M. Lipid composition of human bone marrow. Biochem. J. 95, 252–255 (1965).

31. Dean, J. M. & Lodhi, I. J. Structural and functional roles of ether lipids. Protein Cell 9, 196–206 (2018).

32. Barrachina, M. N. et al. Efficient megakaryopoiesis and platelet production require phospholipid remodeling and PUFA uptake through CD36. *Nat*. Cardiovasc. Res. 2, 746–763 (2023).

33. Paul, S. et al. Shark liver oil supplementation enriches endogenous plasmalogens and reduces markers of dyslipidemia and inflammation. J. Lipid Res. 62, 100092 (2021).

34. Lackner, K. et al. Alterations in ether lipid metabolism and the consequences for the mouse lipidome. Biochim. Biophys. Acta Mol. Cell Biol. Lipids 1868, 159285 (2023).

35. International Mouse Phenotyping Consortium. Mouse phenotype data of knockout mouse lines for protein-coding genes. (2014).

36. Crotty, S. T follicular helper cell differentiation, function, and roles in disease. Immunity 41, 529–542 (2014).

37. Mesin, L., Ersching, J. & Victora, G. D. Germinal center B cell dynamics. Immunity 45, 471–482 (2016).

38. Nojima, T. et al. In-vitro derived germinal centre B cells differentially generate memory B or plasma cells in vivo. Nat. Commun. 2, 465 (2011).

39. Werner, E. R. et al. A novel assay for the introduction of the vinyl ether double bond into plasmalogens using pyrene-labeled substrates. J. Lipid Res. 59, 901–909 (2018).

40. Bozelli, J. C., Jr, Azher, S. & Epand, R. M. Plasmalogens and chronic inflammatory diseases. Front. Physiol. 12, 730829 (2021).

41. Auf dem Brinke, K., et al. Plasma lipidomic patterns associated with disease activity in chronic inflammatory demyelinating polyradiculoneuropathy (LIPID-CIDP). J. Lipid Res. 66, 100903 (2025).

42. Mansell, R. P. et al. Ether lipids influence cancer cell fate by modulating iron uptake. Nat Commun 17, 1835 (2026).

43. Zoeller, R. A. et al. Plasmalogens as endogenous antioxidants: somatic cell mutants reveal the importance of the vinyl ether. Biochem. J. 338 (**Pt 3**), 769–776 (1999).

44. Chen, Z. et al. Ether phospholipids are required for mitochondrial reactive oxygen species homeostasis. Nat. Commun. 14, 2194 (2023).

45. Glaser, P. E. & Gross, R. W. Plasmenylethanolamine facilitates rapid membrane fusion: a stopped-flow kinetic investigation correlating the propensity of a major plasma membrane constituent to adopt an HII phase with its ability to promote membrane fusion. Biochemistry 33, 5805–5812 (1994).

46. Wherley, T. J. et al. Myomaker and ether lipids cooperate to promote fusion-competent membrane states. Cell Rep. 45, 116900 (2026).

47. White, J. K. et al. Genome-wide generation and systematic phenotyping of knockout mice reveals new roles for many genes. Cell 154, 452–464 (2013).

48. Skarnes, W. C. et al. A conditional knockout resource for the genome-wide study of mouse gene function. Nature 474, 337–342 (2011).

49. Pettitt, S. J. et al. Agouti C57BL/6N embryonic stem cells for mouse genetic resources. Nat. Methods 6, 493–495 (2009).

50. Bradley, A. et al. The mammalian gene function resource: the International Knockout Mouse Consortium. Mamm. Genome 23, 580–586 (2012).

51. Schoeler, K. et al. CHK1 dosage in germinal center B cells controls humoral immunity. Cell Death Differ 26, 2551–2567 (2019).

52. Keller, M. A. et al. Studying fatty aldehyde metabolism in living cells with pyrene-labeled compounds. J Lipid Res 53, 1410–1416 (2012).

