## Supplemental material for "Plasmalogen metabolism shapes germinal centre immunity"

### Abstract

Plasmalogens are membrane lipids whose roles in immunity are emerging through identification of the *PEDS1* gene and advances in lipid analytics. Because PEDS1 catalyses the final step of plasmalogen biosynthesis, *Peds1* knockout models plasmalogen loss. Building on reports of a haematopoietic phenotype in *Peds1*-deficient mice, we performed extended profiling to characterise haematological and immune phenotypes. In line with these previous investigations, mice showed reduced red blood cell counts and haematocrit, with increased mean platelet volume. At steady state, plasmalogen deficiency was associated with reduced germinal centre (GC) B cell frequencies in mucosal lymphoid tissues and lower basal serum IgG1 levels. Following immunisation, GC B cell frequencies and ovalbumin-specific IgG1 were similar across genotypes, yet GC B cells exhibited elevated mitochondrial superoxide and plasma cells displayed increased lipid peroxidation. To test for B cell-intrinsic effects, we used an *in vitro* GC culture, where *Peds1*-deficient B cells showed reduced class switching to IgG1, intracellular IgG1 accumulation and diminished IgG1 secretion. Together, these findings support a role for plasmalogens and PEDS1 as a lipid-metabolic modulator of GC dynamics and IgG1 output, consistent with broader ether lipid perturbations. Plasmalogen metabolism may therefore offer a potential entry point to modulate humoral immunity or haematopoiesis.

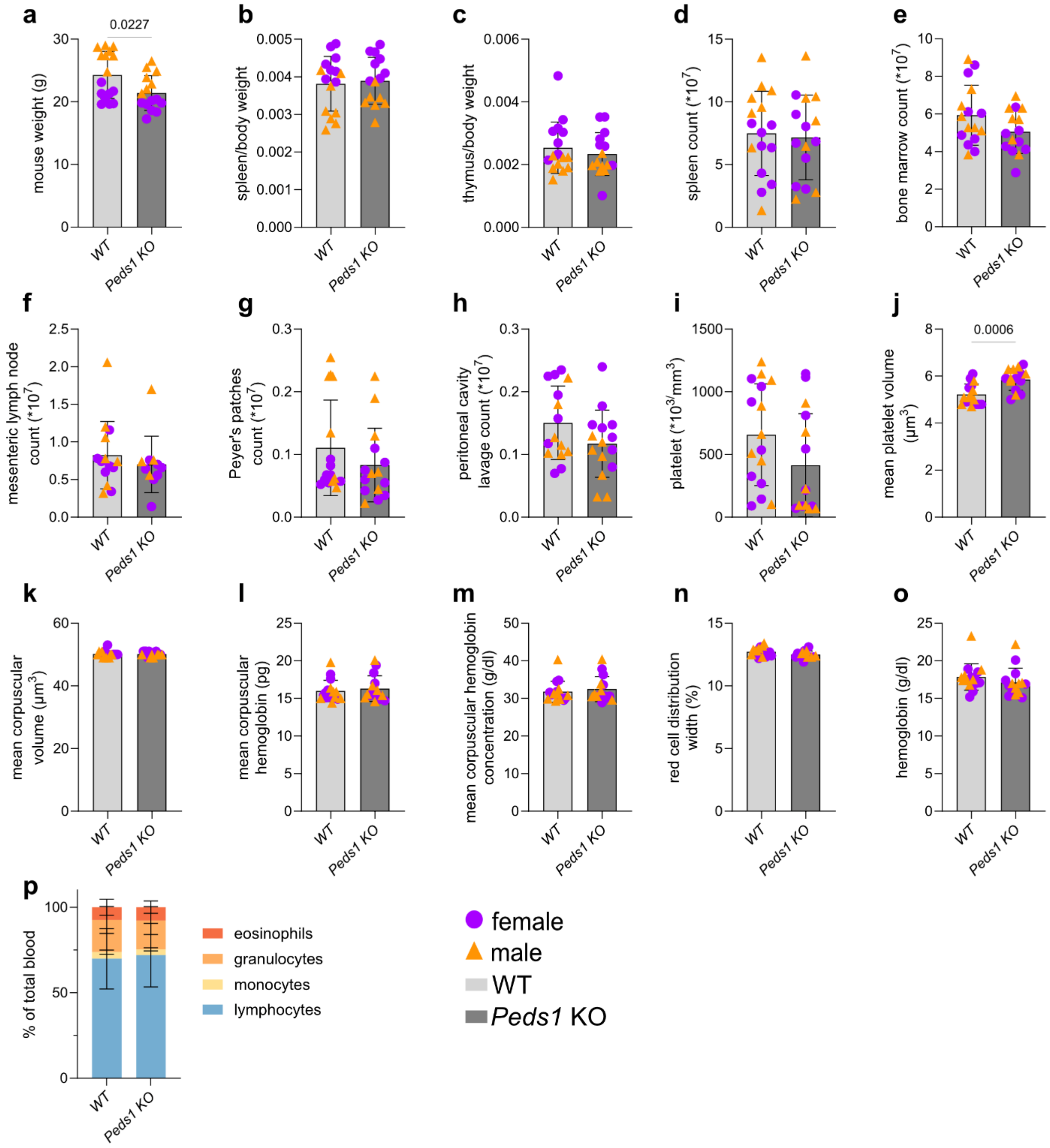


***Supplementary Figure 1: Haematological and organ parameter analysis of WT and* Peds1 *KO mice. a** Mouse body weight. **b** Spleen/body weight ratio. **c** Thymus/body weight ratio. Total cell count of **d** splenocyte, **e** bone marrow, **f** mesenteric lymph node, **g** Peyer's patches, **h** peritoneal cavity lavage. Automated blood counter analysis showing **i** platelet count, **j** mean platelet volume, **k** mean corpuscular volume, **l** mean corpuscular haemoglobin, **m** mean corpuscular haemoglobin concentration, **n** red cell distribution width, **o** haemoglobin concentration and **p** percentage of eosinophils, granulocytes, monocytes, and lymphocytes of total blood leukocytes. WT (n = 16, 8 female, 8 male mice), Peds1 KO mice (n = 15, 8 female, 7 male mice) and females are represented by violet circles and males by yellow triangles. Data are presented as mean ± SD. Statistical comparisons were made using two-tailed Student's t-test (**a-o**) or two-way ANOVA with Bonferroni's multiple comparison test (**p**).

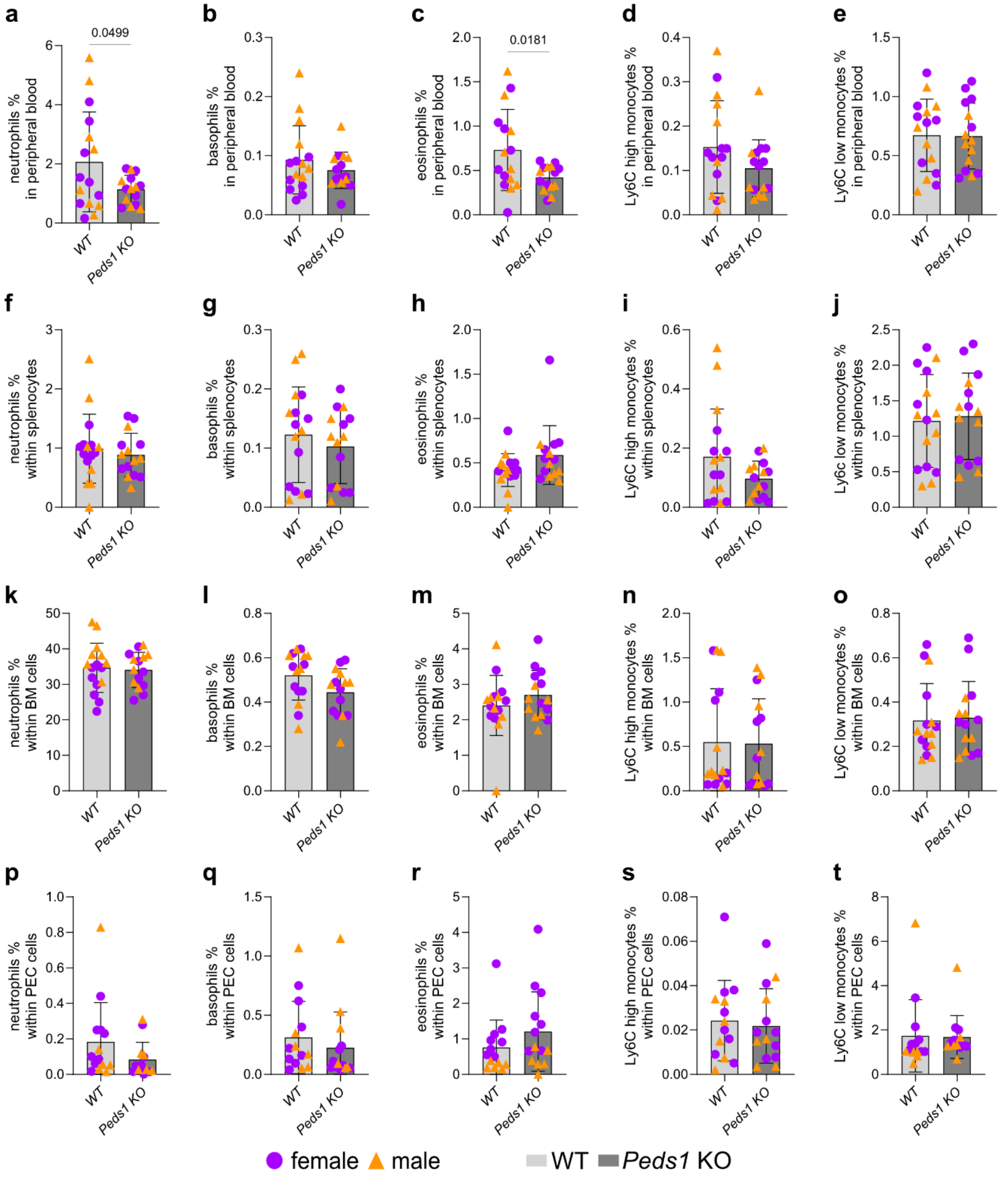


***Supplementary Figure 2: Myeloid cell flow cytometric characterisation of peripheral blood, spleen, bone marrow and peritoneal cavity lavage of WT and* Peds1 *KO mice.*** *Detailed myeloid subsets in PBL:* ***a*** *neutrophils,* ***b*** *eosinophils,* ***c*** *basophils,* ***d*** *Ly6Chigh monocytes,* ***e*** *Ly6Clow monocytes; in spleen* ***f*** *neutrophils,* ***g*** *basophils,* ***h*** *eosinophils,* ***i*** *Ly6Chigh monocytes,* ***j*** *Ly6Clow monocytes; in bone marrow* ***k*** *neutrophils,* ***l*** *basophils,* ***m*** *eosinophils,* ***n*** *Ly6Chigh monocytes,* ***o*** *Ly6Clow monocytes; in peritoneal cavity lavage (PEC)* ***p*** *neutrophils,* ***q*** *basophils,* ***r*** *eosinophils,* ***s*** *Ly6Chigh monocytes,* ***t*** *Ly6Clow monocytes.* ***a-o*** *WT (n = 16, 8 female, 8 male mice),* Peds1 *KO mice (n = 15, 8 female, 7 male mice);* ***p-t*** *for both WT and* Peds1 *KO mice in PEC (n = 16, 8 female, 8 male mice). Females are represented by violet circles and males by yellow triangles. Data are presented as mean ± SD and statistical significance was determined by two-tailed Student's t-test (****a-t****).*

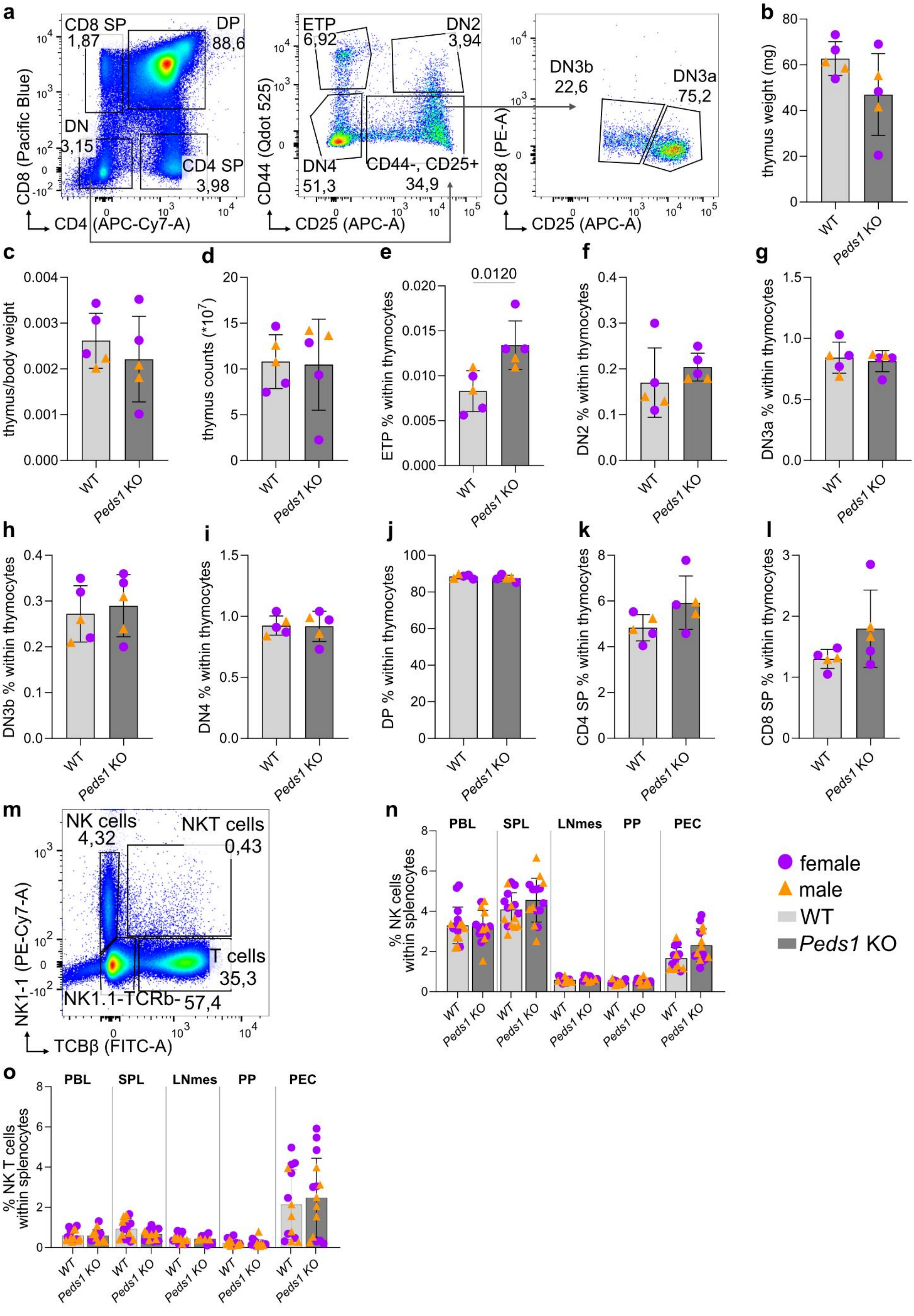

a
CD8 SP 1,87
DP 88,6
DN 3,15
CD4 SP 3,98
CD8 (Pacific Blue)
CD4 (APC-Cy7-A)
ETP 6,92
DN2 3,94
DN4 51,3
CD44-, CD25+ 34,9
CD44 (Qdot 525)
CD25 (APC-A)
DN3b 22,6
DN3a 75,2
CD28 (PE-A)
CD25 (APC-A)
b
thymus weight (mg)
c
thymus/body weight
d
thymus counts (*10^7)
e
0.0120
ETP % within thymocytes
f
DN2 % within thymocytes
g
DN3a % within thymocytes
h
DN3b % within thymocytes
i
DN4 % within thymocytes
j
DP % within thymocytes
k
CD4 SP % within thymocytes
l
CD8 SP % within thymocytes
WT
Peds1 KO
m
NK cells 4,32
NKT cells 0,43
T cells 35,3
NK1.1-TCRb- 57,4
NK1-1 (PE-Cy7-A)
TCBβ (FITC-A)
n
PBL
SPL
LNmes
PP
PEC
% NK cells within splenocytes
female
male
WT
Peds1 KO
o
PBL
SPL
LNmes
PP
PEC
% NK T cells within splenocytes

***Supplementary Figure 3: Thymocytes and splenic NK and NK T cells in WT and* Peds1 *KO mice. a*** *Gating strategy for thymocytes: DN (CD8$^{-}$, CD4$^{-}$), from DN: ETP (CD44$^{+}$, CD25$^{-}$), DN2 (CD44$^{+}$, CD25$^{+}$), from CD44$^{-}$ CD25$^{+}$: DN3b (CD28$^{+}$, CD25$^{-}$), DN3a (CD28−, CD25$^{+}$), DN4 (CD44$^{-}$, CD25$^{-}$), DP (CD8$^{+}$, CD4$^{+}$), CD4 SP (CD8$^{-}$, CD4$^{+}$), CD8 SP (CD8$^{+}$, CD4$^{-}$).* ***b*** *Thymus weight and* ***c*** *thymus/body weight ratio.* ***d*** *Total thymocyte count.* ***e*** *ETP percentage within thymocytes,* ***f*** *DN2,* ***g*** *DN3a,* ***h*** *DN3b,* ***i*** *DN4,* ***j*** *DP,* ***k*** *CD4 SP,* ***l*** *CD8 SP.* ***m*** *Gating strategy for splenic NK cells (TCRβ$^{-}$, NK1.1$^{+}$), NKT cells (TCRβ$^{+}$, NK1.1$^{+}$), and TCRβ$^{+}$ T cells (TCRβ$^{+}$, NK1.1$^{-}$).* ***n*** *NK cells and* ***o*** *NKT cells within PBL, SPL, LNmes, PP and PEC. Thymocytes in WT and* Peds1 *KO mice (n = 5, 3 female, 2 male mice,* ***b-l****) and splenic NK and NK T cells* Peds1 *KO mice (for PBL, SPL, PP, PEC: n = 15, 8 female, 7 male mice, for LNmes: n = 11, 7 female, 4 male mice,* ***n-o****) and WT mice (for PBL, SPL, PP: n = 16, 8 female, 8 male mice, for LNmes: n = 14, 7 female, 7 male mice, for PEC: n = 14, 8 female, 6 male mice,* ***n-o****). Females are represented by violet circles and males by yellow triangles. Data are presented as mean ± SD. Statistical significance was determined by one-way ANOVA with Bonferroni's multiple comparison test.*

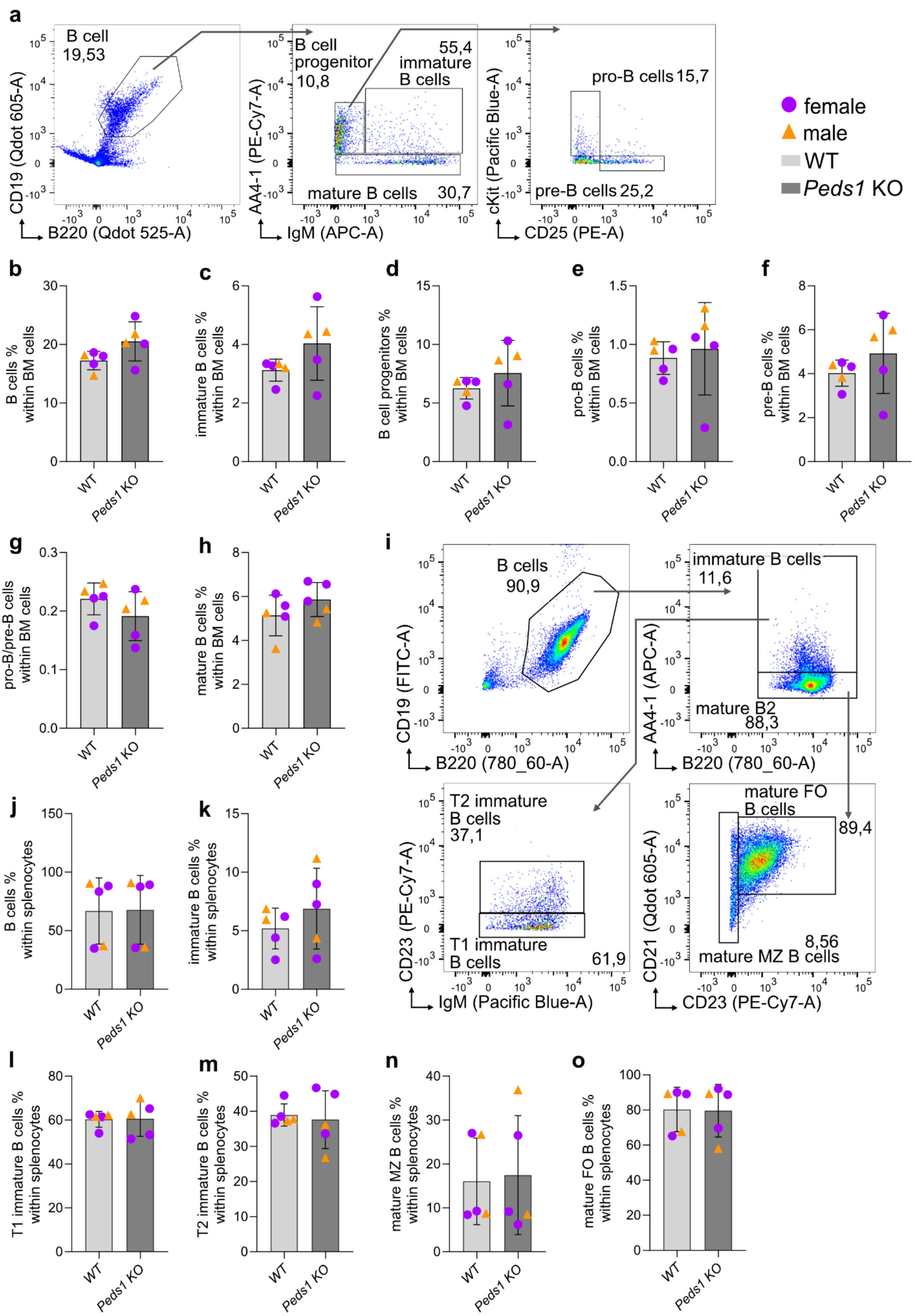
a
B cell
19,53
CD19 (Qdot 605-A)
B220 (Qdot 525-A)
B cell progenitor
10,8
55,4
immature B cells
mature B cells
30,7
AA4-1 (PE-Cy7-A)
IgM (APC-A)
pro-B cells 15,7
pre-B cells 25,2
cKit (Pacific Blue-A)
CD25 (PE-A)
female
male
WT
Peds1 KO
b
B cells % within BM cells
c
immature B cells % within BM cells
d
B cell progenitors % within BM cells
e
pro-B cells % within BM cells
f
pre-B cells % within BM cells
g
pro-B/pre-B cells within BM cells
h
mature B cells % within BM cells
i
B cells
90,9
CD19 (FITC-A)
B220 (780_60-A)
immature B cells
11,6
mature B2
88,3
AA4-1 (APC-A)
B220 (780_60-A)
T2 immature B cells
37,1
T1 immature B cells
61,9
CD23 (PE-Cy7-A)
IgM (Pacific Blue-A)
mature FO B cells
89,4
8,56
mature MZ B cells
CD21 (Qdot 605-A)
CD23 (PE-Cy7-A)
j
B cells % within splenocytes
k
immature B cells % within splenocytes
l
T1 immature B cells % within splenocytes
m
T2 immature B cells % within splenocytes
n
mature MZ B cells % within splenocytes
o
mature FO B cells % within splenocytes
WT
Peds1 KO

***Supplementary Figure 4: Flow cytometric analysis of B cell subsets in WT and* Peds1 *KO mice.*** ***a*** *Gating strategy for pro-B, pre-B, immature IgM+ and mature B cells in the BM.* ***b*** *B cell percentage in BM,* ***c*** *immature B cells,* ***d*** *B cell progenitors,* ***e*** *pro-B cells,* ***f*** *pre-B cells,* ***g*** *pro-/pre-B cell ratio, and* ***h*** *mature B cells.* ***i*** *Gating strategy for splenic B cell subsets. Shown are total $CD19^+B220^+$ B cells, and immature ($AA4.1^+$), T1 ($AA4.1^+$,CD23low, $IgM^+$), T2 ($AA4.1^+$,CD23high $IgM^+$), mature follicular (FO) ($CD21^+$, $CD23^+$), and mature marginal zone (MZ) ($CD21^+$, $CD23^-$) B cells within total $CD19^+B220^+$ B cells.* ***j*** *B cell percentage and* ***k*** *frequency of immature B cells in splenocytes.* ***l*** *T1 immature B cells,* ***m*** *T2 immature B cells,* ***n*** *mature MZ B cells,* ***o*** *mature FO B cells.* Peds1 *KO and WT (n = 5, 3 female, 2 male mice,* ***b-o****)* ***p*** *Percentage of B cells within PEC, WT (n = 14, 8 female, 6 male mice) versus* Peds1 *KO mice (n = 15, 8 female, 7 male mice). Females are depicted by violet circles and males by yellow triangles. Data are presented as mean ± SD. Statistical significance for all B cell percentages was determined using two-tailed unpaired Student's t-test.*

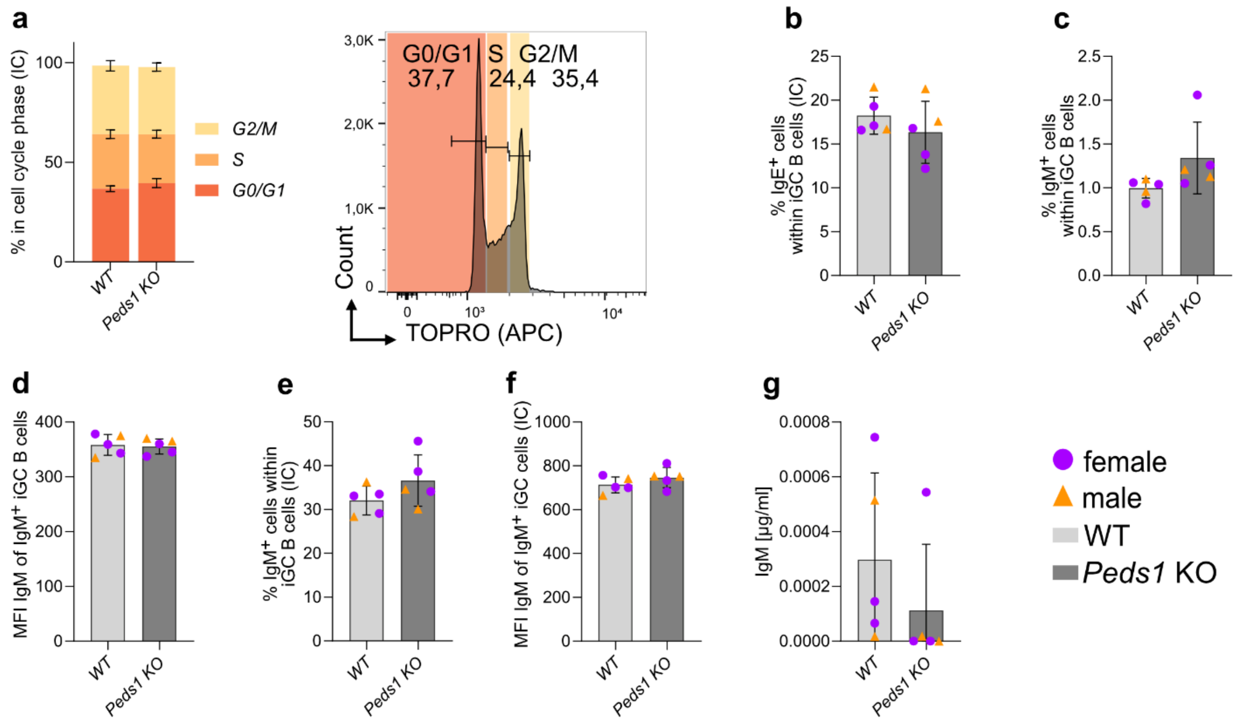


***Supplementary Figure 5:* In vitro *iGC B cell culture analysis of splenic B cells from* Peds1 *KO and WT mice.*** ***a*** *Cell cycle profile of iGC cells after 8 days in culture and representative gating strategy for cell cycle profiling. Fractions of* ***b*** *IgE*$^{+}$ *and* ***c*** *IgM*$^{+}$ *cells within total iGC cells.* ***d*** *IgM MFI of cell surface IgM within iGC B cells.* ***e*** *Fraction of intracellular IgM*$^{+}$ *cells within iGC B cells and* ***f*** *MFI of intracellular IgM within iGC B cells.* ***g*** *Secreted IgM measured in the culture supernatant via ELISA. Data are presented as mean ± SD, for both WT and* Peds1 *KO (n = 5, 3 female and 2 male mice). Females are represented by violet circles and males by yellow triangles. Statistical comparisons were made using two-way ANOVA with Bonferroni's multiple comparison test (****a****) or two-tailed Student's t-test (****b-g****).*

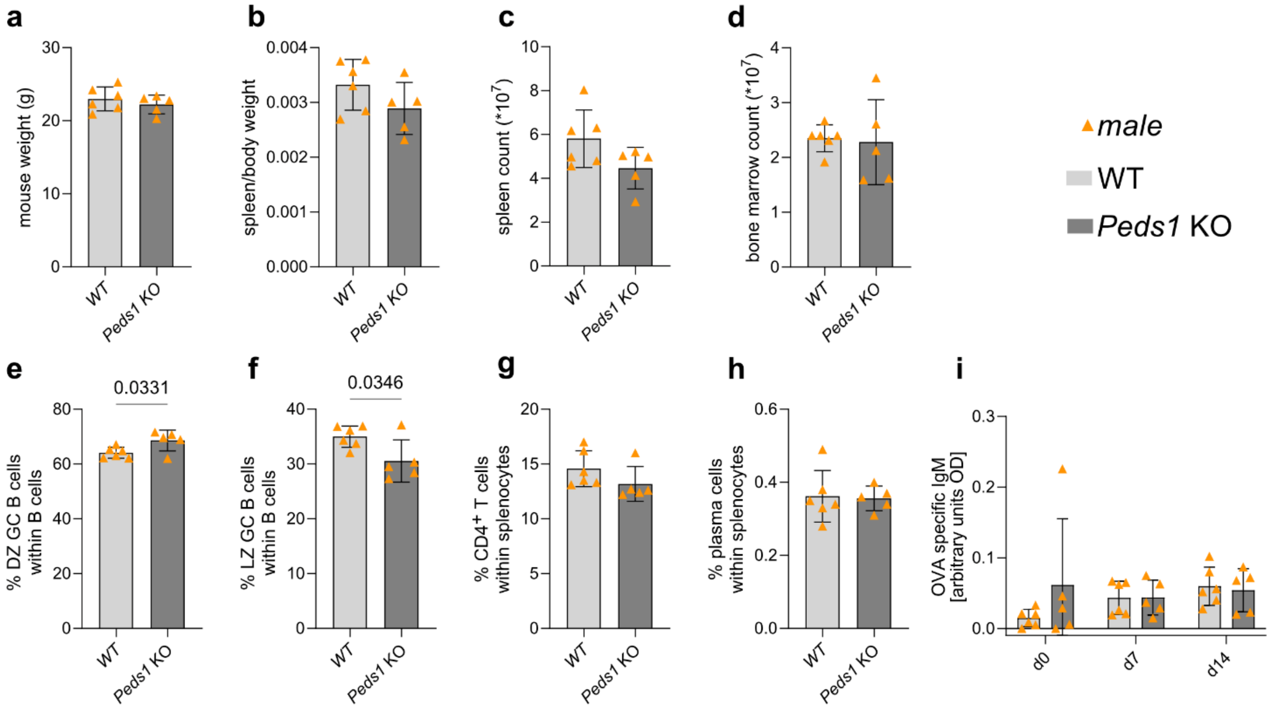


***Supplementary Figure 6: Immune cell analysis of*** **Peds1** ***KO and WT mice at day 14 post OVA immunisation. a*** *Mouse body weight and* ***b*** *spleen/body weight ratio. Total cell counts of* ***c*** *spleen and* ***d*** *bone marrow.* ***e*** *Fractions of DZ (CXCR4$^{high}$, CD86$^{low}$) and* ***f*** *LZ (CXCR4$^{low}$, CD86$^{high}$) cells within B220$^{+}$CD19$^{+}$CD38$^{-}$ GC B cells in spleen.* ***g*** *Fractions of CD4$^{+}$ T cells and* ***h*** *plasma cells (TACI$^{+}$, CD138$^{+}$) among splenocytes.* ***i*** *OVA-specific IgM in serum, measured in arbitrary units (optical density, OD). Each data point represents one individual male WT (n = 6) or male* Peds1 *KO (n = 5)* mouse*. Data are shown as mean ± SD. Statistical comparisons were made using two-tailed Student's t-test (****a-h****) or two-way ANOVA with Bonferroni's multiple comparison test (****i****).*
